# Defect-Engineered Metal-Organic Frameworks as Nanocarriers for Pharmacotherapy: Insights into Intracellular Dynamics at The Single Particle Level

**DOI:** 10.1101/2024.04.19.590224

**Authors:** Ge Huang, Marcus Winther Dreisler, Jacob Kæstel-Hansen, Annette Juma Nielsen, Min Zhang, Nikos S. Hatzakis

## Abstract

NanoMOFs are widely implemented in a host of assays involving drug delivery, biosensing catalysis, and bioimaging. Despite their wide use, the cell entry pathways and cell fate remain poorly understood. Here we have synthesized a new fluorescent nanoMOF integrating ATTO 655 into surface defects of colloidal nano UiO-66 that allowed us to track the spatiotemporal localization of Single nanoMOF in live cells. Density Functional Theory(DFT) reveals the stronger binding of ATTO 655 to the uncoordinated saturated Zr_6_ cluster nodes compared with phosphate and Alendronate Sodium (AL). Parallelized tracking of the spatiotemporal localization of tens of thousands of nanoMOFs and analysis using machine learning platforms revealed whether nanoMOFs remain outside as well as their cellular internalization pathways. To quantitatively assess their colocalization with endo/lysosomal compartments, we developed a colocalization proxy approach relying on the nanoMOF detection of particles in one channel to the signal in the corresponding endo/lysosomal compartments channel, considering signal vs local background intensity ratio (S/B) and signal-to-noise ratio (SNR). This strategy effectively mitigates the potential inflation of colocalization values arising from the heightened expression of signals originating from endo/lysosomal compartments, it also overcomes limitations of low SNRs in the endo/lysosomal compartments marker channel, which incapacitates any trajectory-trajectory colocalization assessment. The results accurately measure the amount of nanoMOFs’ colocalization in real-time from early (EE) to late endosomes(LE) and lysosomes(LY) and emphasize the importance of understanding their intracellular dynamics based on single-particle tracking (SPT) for optimal and safe drug delivery.

## 1. Introduction

Metal–organic frameworks (MOFs) are a class of compounds consisting of metal ions and organic ligands, a subclass of coordination polymer. Based on their large porosity, high surface areas, and tunable structures^[1]^, these porous materials are at the forefront of cutting-edge research, drug delivery systems, and biosensing. The nearly infinite range of metal ions and ligands available to form MOF structures^[2]^ allows almost infinite tunability. Despite their potential, it is technically hard to achieve a suitable carrier based on MOFs, that would also be suitable for bioimaging applications. The key requirements include chemical and photostability, small size (on the scale of subcellular structures), biocompatibility, and brightness as well as being soluble in the aqueous, reactive, and crowded biological environment. Some of these can be overcome by utilizing common coordination groups (–COOH in most cases) that usually possess a small p-orbital conjugation, which leads to a large energy band gap (Eg) for a vast majority of MOFs such as MOF-5 (Eg = 3.4 eV), ZIF-8 (Eg = 5.5 eV), MIL-125 (Eg = 3.6 eV)^[3]^.

The MOFs’ physical properties like particle size at the micro level or millimeter level, on the other hand, challenge the applications of cell imaging and cell uptake limiting our understanding of their interaction with cellular membranes. A quantitative understanding of the interaction between nanoMOFs and cell membranes, the cell uptake mechanisms, and intracellular localization in drug delivery systems are essential for safe and efficient therapeutic applications of MOFs^[4]^.

Here, we addressed the above challenges and developed a colloidally stable defect-engineered multivariate (MTV) MOF drug carrier and directly imaged its internalization pathways in cells. We tailored nanoMOF characteristics by tuning the temperature synthesis parameters that control MOFs physical properties such as size, crystallinity, colloidal dispersion, stability, and porosity^[5]^. Using a low temperature-induced defect formation strategy^[6]^ we found a positive correlation between the volume defects and surface defects, the defects based on the number of missing linkers caused by uncoordinated saturated Zr_6_ clusters (Figure 1a). We characterized the relationship between their crystal growth, porosity, missing ligands analysis, and visualization of surface defects. The MTV MOF platform was then utilized as a drug delivery system, adopting an inside (doped phosphate anti-cancer drug AL)-outside(doped carboxyl ATTO 655 fluorescence molecular) approach for post-synthesis modification (Figure 1b).

**Figure 1:**
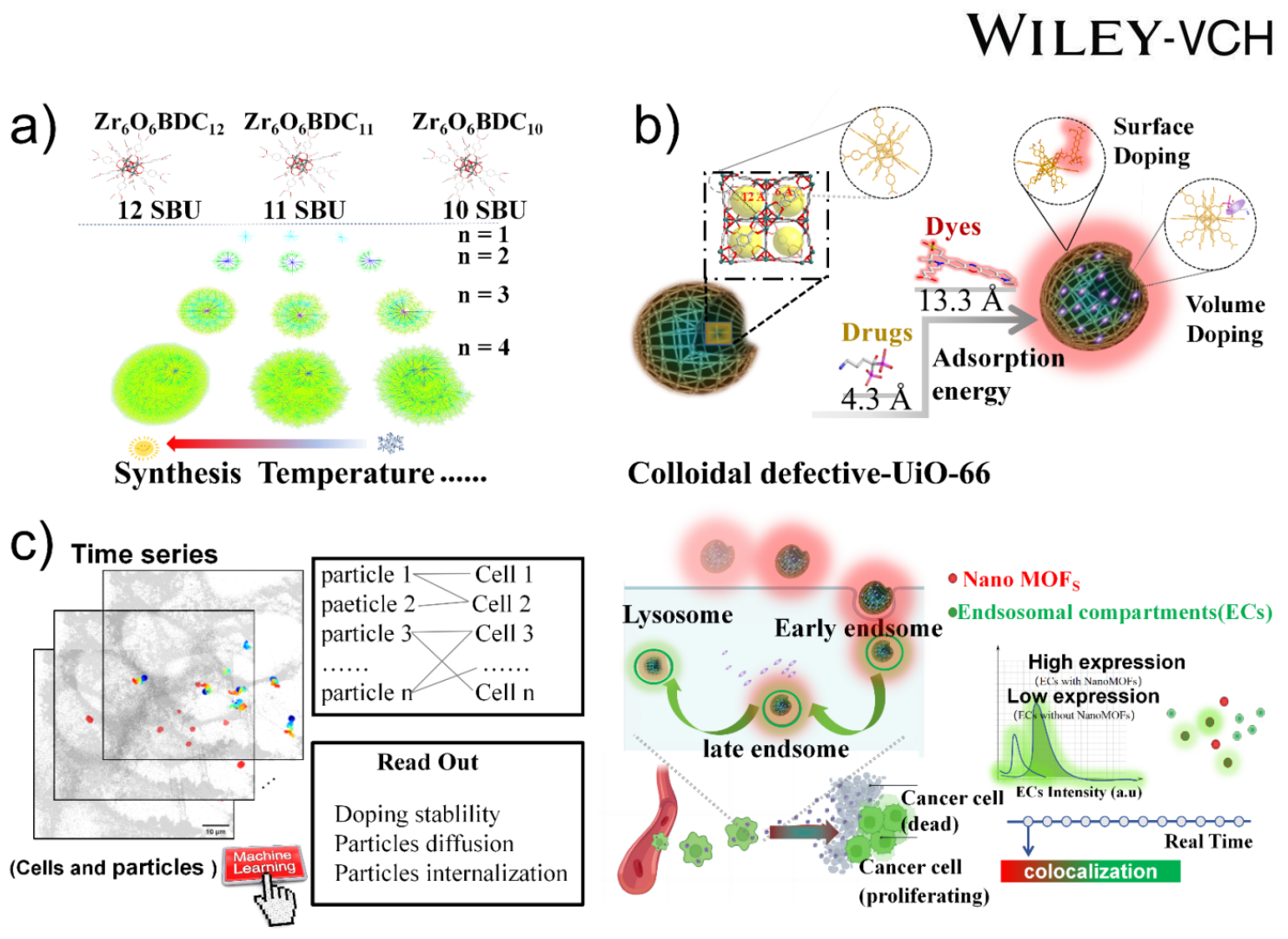
Representation of the methodology development of nanoMOFs as drug delivery vehicles. a) Cartoon representation of missing linker’s dynamic growth in UiO-66 by 12, 11 and 10 branches of tree diagram. b) Schematic diagram of surface modification and volume modification. c) Representation of the framework for parallelized tracking of cell entry pathway of nanoMOFs in HeLa cells aided by machine learning analysis (left); Proposed cell uptake mechanism of ATTO-UiO-66@Al based on the tracking data and the algorithm for colocalization percentage between endo/lysosomal compartments and nanoMOFs (right).

Conventional characterization techniques of MOF interaction with cells primarily report the ensemble average value of a large number of particles, often masking the heterogeneous behavior of particles in complex biological environments. Single-particle tracking (SPT)^[7,8,9,10]^ techniques, on the other hand, offer the direct observation of the motion of individual particles and combined with sophisticated analysis of the heterogenous motion have successfully addressed multiple basic biological problems successfully. Capitalizing on this toolbox allowed us to observe the diverse and heterogeneous nanoMOFs internalization details in live HeLa cells. To account for the challenges associated with fast moving particle at dense environments and with low signal to noise, we developed a colocalization proxy approach relying on the detection of particles in one channel (nanoMOFs) and subsequent consideration of signal in the corresponding endo/lysosomal compartments channel. This advanced approach to tracking nanoMOFs provides critical insights into their intracellular dynamics and interactions, crucial for enhancing their therapeutic efficacy (Figure 1c).

## 2. Results and Discussion

### 2.1 Defect regulation synthesis of colloidal defective UiO-66 for molecular doping in the particles

To optimize MOFs as drug delivery systems via defect chemistry (Figure 2a), the linear correlation in the synthesis temperature, N_2_ uptake, TGA, dye surface modified amount, and contact angle were analyzed (Figure S1-S9). The Pearson value in each two variables is mostly between 0.82 and 0.99 (Figure 2b and Table 1 and Table 2). This is consistent with the hypothesis that a lower synthesis temperature results in a more linker deficient and hence more porous framework and more uncoordinated saturated metal site on the surface of the crystals^[1c, 5]^.

**Figure 2:**
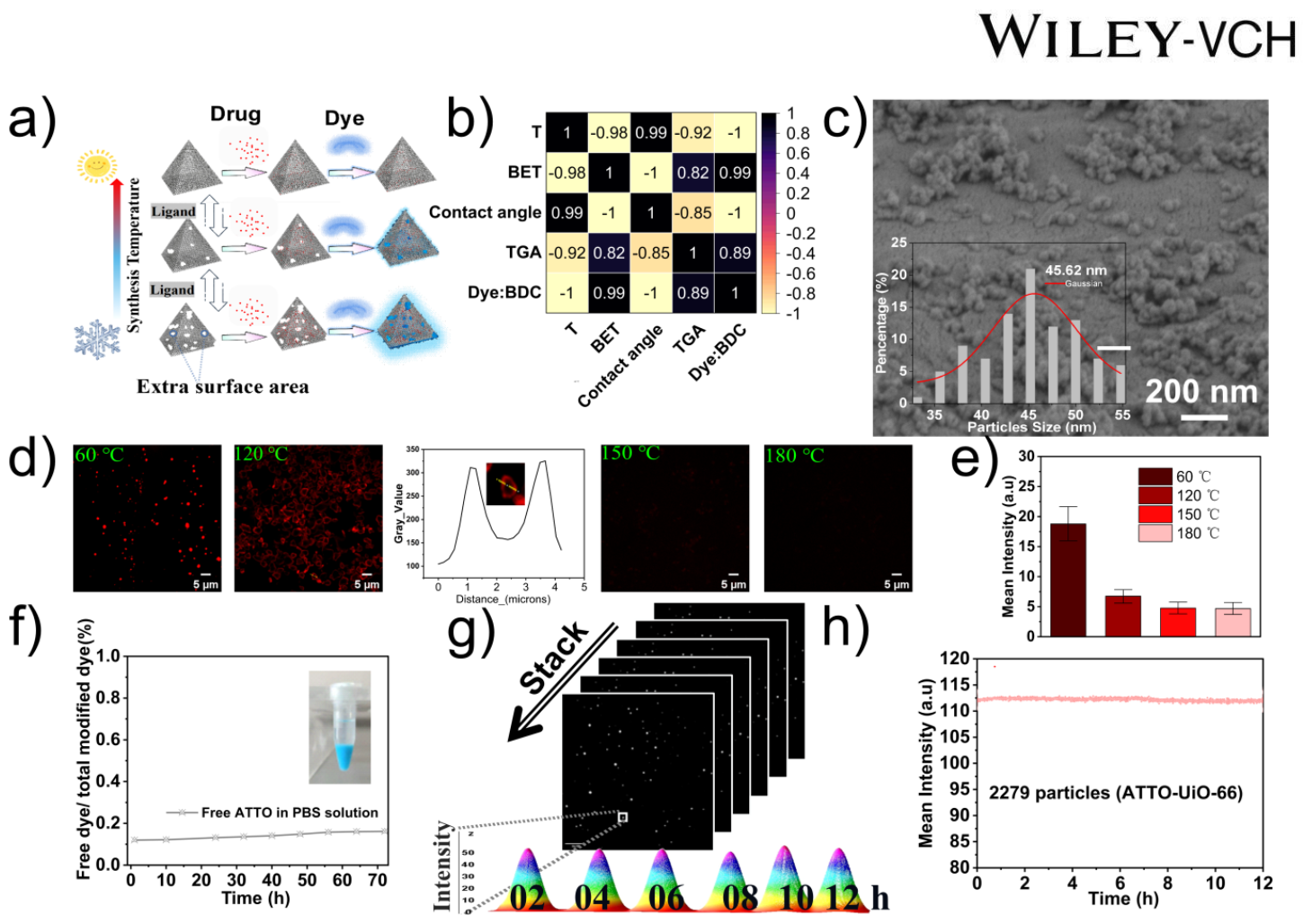
Characterization of nanoMOFs as a dye and drug-doped nanocarrier based on defect chemistry. a) Cartoon showing drug loading and dye modification based on MOF’s defect engineering, b) The correlation plot shows a positive correlation between surface defects and volume defects compared to the synthesis temperature T (°C), BET (cm^3^/g), contact angle (°), TGA (ZrO_2_ %), and the rate of dye/BDC (mol/mol). Positive values mean positive correlation from 0 to 1, and negative values mean negative correlation from -1 to 0. c) Representative SEM image of ATTO-UiO-66 displaying the relative monodisperse dimensions (average size = 45.62 nm). d) Spinning disc microscopy images of ATTO 655 signal of the surface-immobilized ATTO-UiO-66 prepared at conditions resulting in diverse defect densities (green: synthesis temperature). e) The mean intensity of different defect levels UiO-66 immobilized by ATTO 655. f) Monitor signal of leakage of ATTO 655 from ATTO-UiO-66 in upper liquid for 3 days (see supporting information S15) (Inside: A picture of 5 mg ATTO-UiO-66 is stable in 1 mL PBS solution for 30 days). g) Evaluation of photostability by recordings of individual ATTO-UiO-66 for 12 h (30% Laser 640 power, exposed time 50.04 ms and 30 s per frame). h) The mean intensity of 2279 ATTO-UiO-66 particles for 12 h. (error bar per frame based on n = 2279 particles standard deviation per frame).

To verify the chemical modification of the dye on the surface of the MOF crystal, we washed 5 times with the deionized water and until no UV-Vis adsorption of fluorophores was detected in the supernatant. Then, we used FTIR to analyze two samples with the same mass ratio for the host (defective UiO-66) and guest (Gallocyanine): one sample underwent chemical modification and the other sample was a physical mixture based on NMR results (Figure S10). We found that the fluorophores Gallocyanine is modified on the crystals in the state of deprotonation, supporting that the carboxyl dye is bonded with these metal sites with the hanging bond. Confocal Fluorescence Microscopy, identified linear areas well above the diffraction limit not loaded with fluorophores supporting the presence of defects on the outer surface of UiO-66 crystals (Figure S11) colocalized with the carboxyl gallocyanine. Such features could be related to the growth mechanism of the UiO-66 crystal surface, as has been previously suggested for liposomes or other coordination compounds^[1c,11]^. This colocalization is consistent with positive correlation results of missing links’ amount between volume defects and surface defects (Figure 2b).

As monodisperse, colloidally stable samples are imperative for drug delivery, we continued to decrease the synthesis temperature to 60 °C. According to our previous work report^[6]^, we can synthesize many defective UiO-topology nanoparticles like UiO-66, UiO-67, DUT-52, and PCN-56. Complicating its detection in live cell studies due to spectral overlap with most common dyes used for labeling of cellular membranes and components, we chose ATTO 655 for surface modification. The SEM image shows that the ATTO-UiO-66 particle size is about 45.62 nm (Figure 2c). Representative confocal figures and line scan in Figure 2d supports the fluorescence attachment is on the defective MOFs (Figure 2d). In contrast, no fluorescence is observed on near defect-free (180 °C) MOFs (Figure 2d). These results are consistent with the NMR results of three different samples with dye modification (Figure S9, Table1).

The uncoordinated saturated metal sites present within the internal defects (volume defects) of the Metal-Organic Framework (MOF) facilitate the high loading of phosphate drugs Alendronate Sodium due to the affinity between the MOF structure and phosphorus atoms^[12]^. Externally, the uncoordinated sites are modified with ATTO 655, The PXRD results show that the MOF retains its core structure even after the loading of drugs and the modification with fluorescent molecules on its surface (Figure S12). Interestingly, due to the high drug loading and the involvement of phosphonate groups in coordination, upon loading AL, a new diffraction peak appears around 10° in the PXRD (Figure S12). DFT calculations show that ATTO 655 strongly doped in defect UiO-66 surface chemically (Figure S13). This modification preference arises because MOFs have a higher affinity for the Alendronate Sodium (AL) pharmaceutics compound as compared to ATTO 655 based on adsorption energy (Table 3). To verify the loading of the drug we compared the supernatant drug Abs intensity (at wavelength = 205 nm) before and after loading drugs into the MOFs with synthesis temperature (60, 120, 150, 180 °C) (Figure S14 and S15). This showed it to be 28% AL for the nanoMOF (synthesized at 60 °C) and 30% for powdered nanoUiO-66@AL analyzed by ICP (Zr : P).

The formation of stable Zr−O−P bonds at the structurally uncoordinated missing linker site was shown by some reported works: Guan et al. showed that various functionalized UiO-66 MOFs removed phosphate from acidic solvents^[13]^. Using a Zr-based MOF, Wang et al. demonstrated that phosphate adsorption occurred at a missing linker site via linker exchange^[14]^. These experiments clearly demonstrate that the moieties connected to the six metal atoms of the SBU influence P adsorption. The size of the drug particles (4.3 Å) is smaller than the windows of UiO-66 (6 - 12 Å) suggesting that we loaded the phosphates drug (AL) in the volume defects.

The surface defects play a role in modifying the fluorescence molecule ATTO 655 (13.3 Å)^[15]^. After the synthesis of ATTO-UiO-66 for luminescent MOFs^[16]^, the stability of this hanging bond was tested under the PBS solution at 37 °C for 72 hours by monitoring the fluorescence intensity of dye at the excitation wavelength at 600 nm in the upper liquid at 1, 10, 24, 32, 40, 48, 54, 64, 72 h (Figure 2f and Figure S16). The results show that the concentration of dyes in the upper liquid didn’t increase after 3 days supporting its stable association. Treatment with acetic acid at day 3, resulted in increase of fluorescent signal on the supernatant consistent with the dye being separated from the surface of UiO-66. Therefore these hanging bonds through the defect chemistry had a high stability for at least 3 days in PBS solution.

To evaluate further the irreversible association of ATTO 655 as well as the photostability of ATTO-UiO-66. we performed single particle readouts. Glass surface passivated with positively charged poly-L-lysine surfaces were used to immobilize ATTO-UiO-66 (zeta potential: -7.14 mV) in the growth medium. ATTO 655 labeled UiO-66 was added on the poly-L-lysine surface without diffusion on the surface and their photo-stability was observed by spinning disk confocal microscopy for 12 hours (Figure 2g and Movie 1). Extraction of the background corrected signal of the particles, confirmed the mean intensity of ATTO-UiO-66 to be stable for 12 h supporting both the stable association of ATTO 655 and the photostability of ATTO-UiO-66 for live cell imaging (Figure 2h).

### 2.2 Internalization of nanoMOFs for Drug Delivery

NanoMOFs have been reported to be delivered in cells; however, their internalization mechanism remains unclear due to the absence of direct observations. To resolve this, we recorded their interaction with cell membranes and their internalization and trafficking in HeLa cells through single-particle tracking (SPT)^[7,8,9,10]^ studies. Particle localization, intensity calibration, and image analysis were based on our image analysis pipelines and our machine-learning toolboxes^[7a,8]^. Our direct recordings enabled the observation of the spatiotemporal localization of hundreds to thousands of nanoMOFs simultaneously, providing robust quantitative analysis of the spatiotemporal trajectories and revealing the heterogeneous diffusional properties^[9]^ masked by ensemble studies. The recorded heterogeneities in diffusion that can be attributed to nanoparticle and cell interactions, allowing us to classify nanoparticle trajectories as inside or outside a cell or moving on the membrane of the cell.

The diffusional behavior of nanoparticles depends on their interaction with the cellular membrane. We utilized existing machine-learning toolboxes to quantify the interaction between nanoMOFs and cells^[10]^. The experimental and analytical pipeline is outlined in Figure 3a: 1, Two-color parallel recordings of nanoparticles and cells; 2, Cell segmentation and boundary identifications using cellpose^[17]^; 3, Identification of centroids using scikit-image; 4, Tracking of nano MOF and post-processing exceeding a minimum track duration. Final tracks are evaluated to extract the fraction of particles internalized, on the membrane, in solution, or a combination of the above. We then quantified the ratio of the ATTO-UiO-66 particles that are cell internalized vs. the particles in solution for incubation times of 4 h, 8 h, and 24 h. While there appears to be a positive trend, the large variation across experimental conditions challenges the conclusion on the time-dependent increase in internalization, but all data clearly support that the ATTO-UiO-66 particles are indeed internalized (Figure 3b, Figure 3c, and Table 4). Consistent with these, Z projection of the volumetric cell images shows that while some particles diffuse on the extracellular matrix, the majority of particles are indeed internalized by HeLa cells (Figure 3d).

**Figure 3:**
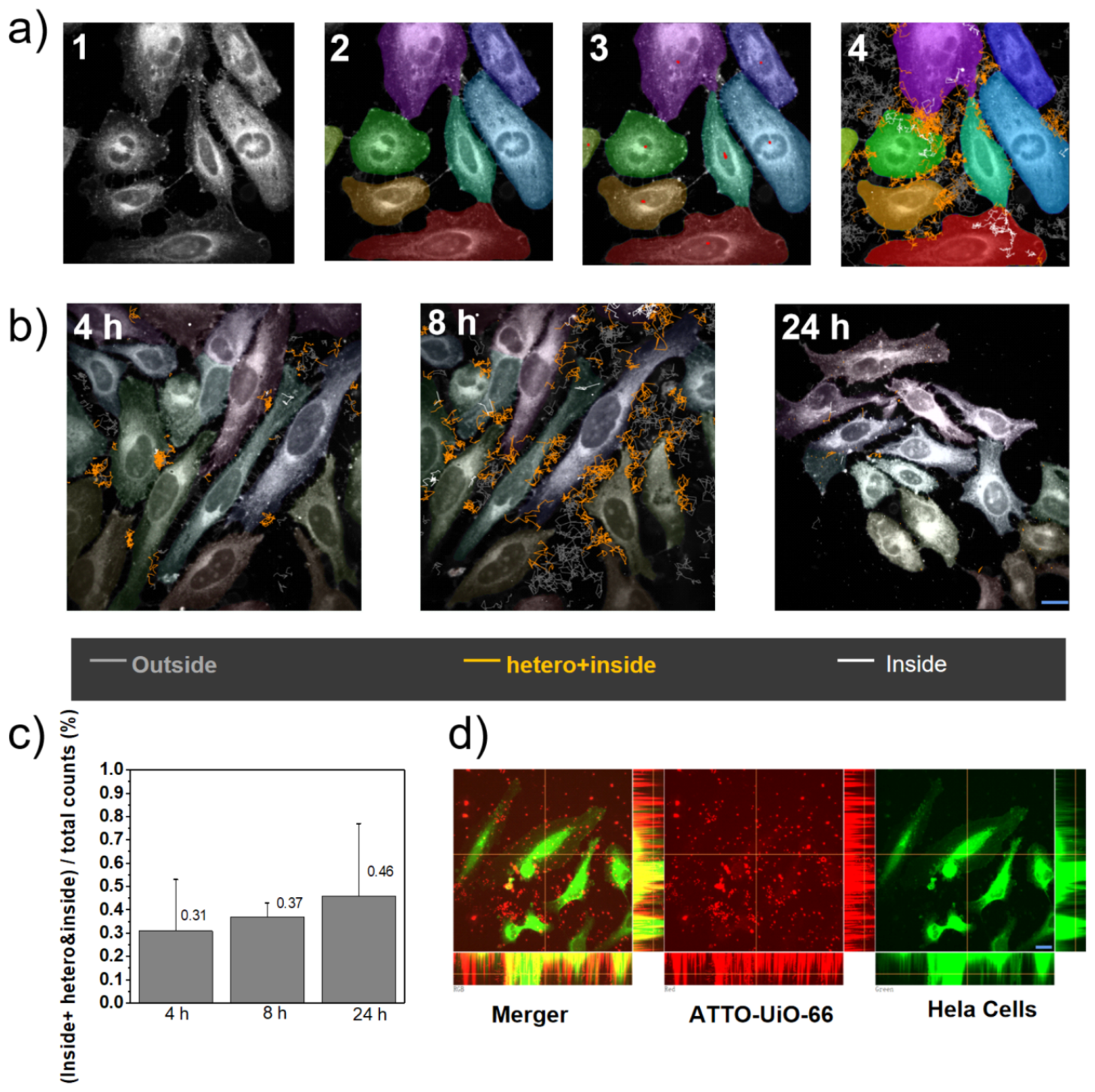
Direct observation of the spatiotemporal localization of individual nanoMOFs internalized in HeLa cells by single-particle tracking (SPT)^[7,8,9,10]^. a) Cell segmentation based on a machine learning algorithm: 1. Finding the frame 0 in live cell stacks, 2. Cell segmentation using cellpose, 3. Finding particle centroids using scikit-image, 4. Tracking and linking centroids using trackpy and post-processing by requiring a minimum track duration. (Scale bar = 20 μm). b) Representative single-plane images acquired by spinning disc microscopy taken at 4 h, 8 h, and 24 h displaying the increased density of particles close to the cells and their internalization. Cells are segmented using cellpose and are displayed by diverse colors, gray trajectories (outside the cells), orange trajectories (some part is on the cell membrane and some part is inside the cells), white trajectories (inside the cells) (Scale bar = 20 μm). c) NanoMOFs are internalized in cells as shown by the internalization percentage with the incubation time. Data analyzed by the machine learning toolbox in Figure 4a. Error bars based on 3 independent experiments. d) Z-stage image stack shows a 3D image of ATTO-UiO-66 inside HeLa cells acquired by spinning disc microscopy (HeLa cells: green channel; ATTO-UiO-66: red channel) (Scale bar = 20 μm).

We then evaluated the internalization pathway of NanoMOFs. Figure 4a shows a schematic diagram of a potential internalization pathway from early endosomal uptake to a potential final destination in lysosomes^[18]^. To assess their pathway, we used ATTO-UiO-66 and transient expression of the GFP-tagged endosomal markers Rab 5 and Rab 7 for early (Figure 4b) and late (Figure 4c) endosomes, respectively^[19]^. Additionally, co-localization with lysosomes (Figure 4d), stained using Lysotracker^[20]^, predominantly occurs in the perinuclear region, signifying the eventual translocation of ATTO-UiO-66 nanoparticles to lysosomes (Movie 2, 3, 4). Using single-particle tracking (SPT)^[7,8,9,10]^, each particles’ trajectory can be recorded (Figure S17), and step length and mean step length analysis were extracted for nanoMOF particles in early endosomes, late endosomes, and lysosomes (Figure 4e).

**Figure 4:**
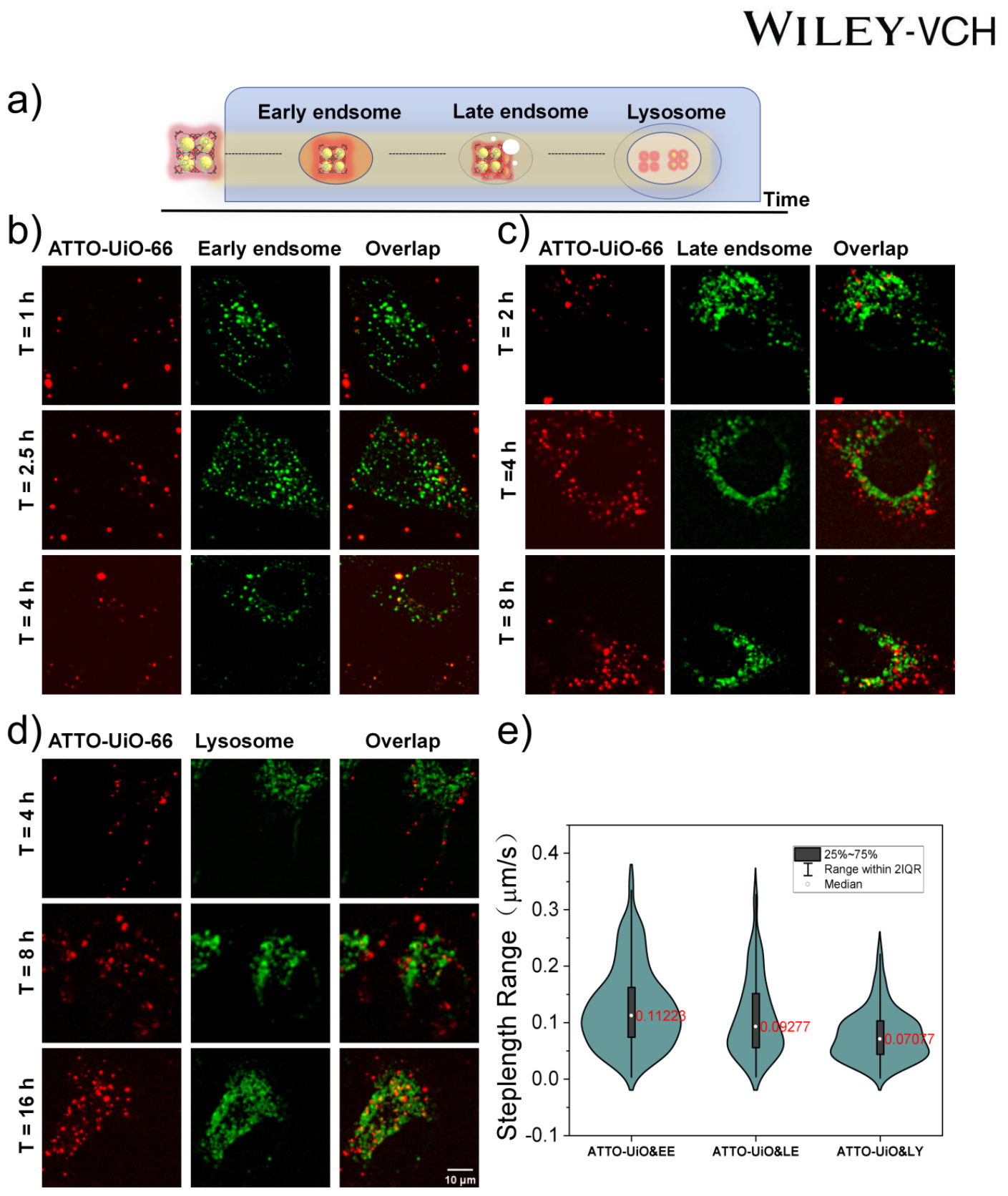
Direct observation of intracellular trafficking of ATTO-UiO-66. a) Schematic showing intracellular trafficking of ATTO-UiO-66. Spinning disk fluorescence microscopy images showing co-localization of ATTO-UiO-66 (640 nm laser, red channel) with b) early endosomes (488 nm laser, green channel) labeled with green fluorescent protein GFP Rab 5. c) late endosomes (green channel) labeled with expression of GFP Rab 7. d) lysosomes (green channel) labeled with LysoTracker® Green (Scale bar = 10 μm). e) The step length range of ATTO-UiO-66 in the early endosome (n = 199), late endosome (n = 113), and lysosome (n = 197) (marked the mean (red) step length).

To overcome the limitations of low SNRs in the endosomal/lysosomal marker channel, which challenges any trajectory-trajectory colocalization assessment, we developed a colocalization proxy approach relying on the detection of particles in one channel (nanoMOFs) and subsequent consideration of signal in the corresponding channel (endo/lyso) at nanoMOFs’ locations (Figure 5a and 5b), The detected nanoMOFs are used as an origin to assess a potential particle-like signal in the endo/lyso channel (Figure 5c). To probe if there exists a particle with its point spread function, and to correct for background, a donut geometry with radii 2 px (1 px = 183 nm) and 5 px was used to define (inner area) a local background at the tail of the point spread function (see methods in support information). The endo/lyso markers were taken as particles and hence colocalized with nanoMOFs (since nanoMOFs defined their origin) if both S/B_d > 1.1 and SNR > 1.1. It is possible to consider colocalization both on a particle basis (mean by ID) and a detection basis. NanoMOFs could colocalize with endo/lyso markers in parts of their trajectory, all of the trajectory, or not at all (Figure S18). We chose to assess colocalization on a detection basis to rightfully include nanoMOFs that colocalized in parts of their trajectory. Considering colocalization on a particle basis (Figure 5d) while still observing high mean signal multipliers S/B_d and SNR further supports the association of nanoMOFs to endo/lyso markers. The reported percentages of colocalization were calculated from the mean and standard deviation in the three biological replicates (Figure 5e, S19, S20, S21, S22, S23) - In two instances, there were just two biological replicates (see supporting information Table 5 for specifications).

**Figure 5:**
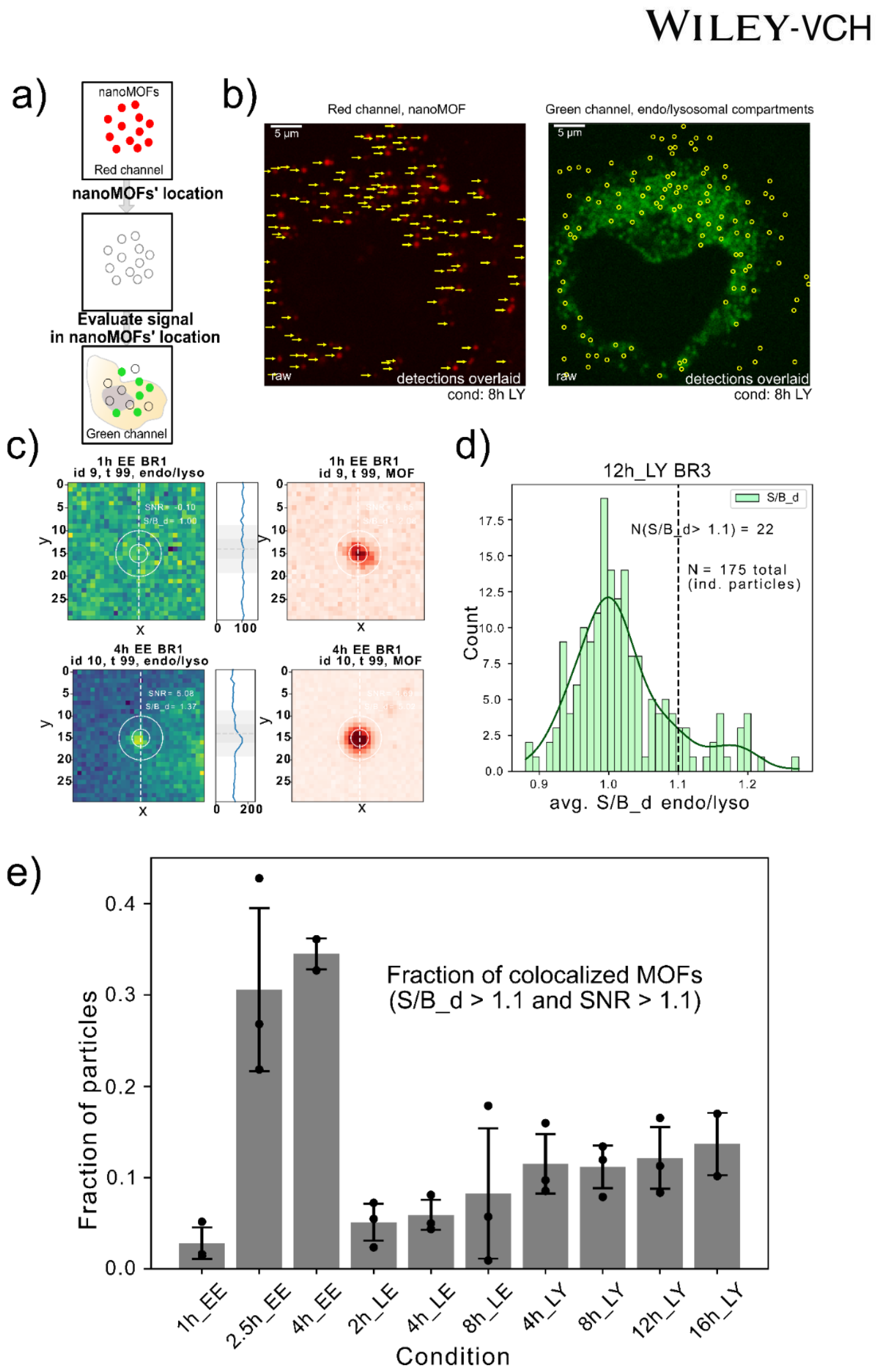
Colocalization analysis of internalized nanoMOFs and endo/lysosomal compartments by single-particle tracking (SPT)^[7,8,9,10]^ in HeLa cells. a) Colocalization algorithm assessing particle-like signal in the endolysosomal channel at nanoMOF locations. The primary steps in the approach are: 1, detecting particles in the nanoMOF channel. 2 identifying nanoMOFs’ locations in endo/lysosomal compartments channel. 3, evaluating signal vs local background intensity ratio and SNR ratio in geometric donuts (1 pixel = 183 nm, radii R1 = 2 px, R2 = 5 px). b) Two channels of a raw single-cell image of a biological replicate in the 8h LY condition. Left: The red nanoMOF channel with arrows depicting where nanoMOFs are detected. Right: The endo/lysosomal marker channel with nanoMOF locations marked c) A 30 x 30 pixels crop-out (centered around nanoMOFs) of the two channels from the 1h EE and 4h EE condition (1 pixel = 183 nm). The detected nanoMOFs (right images) is used as an origin to assess a potential particle-like signal in the endo/lyso channel. A colocalized and a non colocalized area are depicted. d) Histogram of the average S/B_d ratios by particle in the endo/lyso channel from the 12 h LY biological replicate no. 3 (BR3) condition. Kernel Density Estimation (KDE) plotted on top suggests a bimodal distribution. 12.5 % (22/175) of MOFs seem to colocalize with the endo/lyso channel when applying a conservative threshold of 1.1 on the average S/B_d. e) Fraction of colocalized particles by condition calculated on a detection basis by assessing how many detections have SNR > 1.1 and S/B_d > 1.1. Reported fractions and their errors are taken as the mean and standard deviation of three biological replicates.

Figure 5e shows that the colocalization with early endosomes starts at around baseline at 1 hour, increases slightly at 2.5 hours, and stabilizes at 4 hours. The colocalization with late endosomes seems to be consistently above the baseline at all measured time points (2, 4, and 8 hours), suggesting a stable interaction or presence of nanoMOFs in the late endosomes over time. Similarly, a relatively stable colocalization with lysosomes from 4 to 16 hours is observed within error (Figure 5e). In conclusion, these data suggests that nanoMOFs show varying degrees of colocalization with different cellular structures over time, with a notable increase in early endosomal colocalization as time progresses. This could reflect the cellular trafficking pathway of nanoMOFs, from early endosomes to lysosomes.

We then examined whether the defect-engineered nanoMOFs can be effectively utilized for the delivery and release of cargo in HeLa cells^[21]^. We evaluated the loading and release of the anticancer drug AL by quantifying the amount released from nanoMOFs over time (Figure 6a). The assessment of AL release from UiO-66 NPs (28 wt % AL loading) was conducted at 37 °C in the PBS buffer at pH values of 5.0 and 7.4 by monitoring UV-Vis absorption at 205 nm based on the linear relation between Abs and drug concentration (Figure S14, S15). AL release was practically linear for both pH values (see method in the supporting information). 45 % of AL was released from UiO-66 NPs within 50 h at pH 7.4, whereas the release appears slightly increased (58 %) at the same time interval at pH 5.0. The small potential pH-sensitive release trait is linked to phosphates becoming protonated in acidic environments, thereby diminishing the interaction potency between AL and Zr-O clusters in UiO-66. This slightly increased rate of release for acidic pH could be useful as it could impede premature drug release from UiO-66 nanocarriers during circulation while enhancing intracellular drug release in acidic environments found in endosomes or tumor cells.

**Figure 6:**
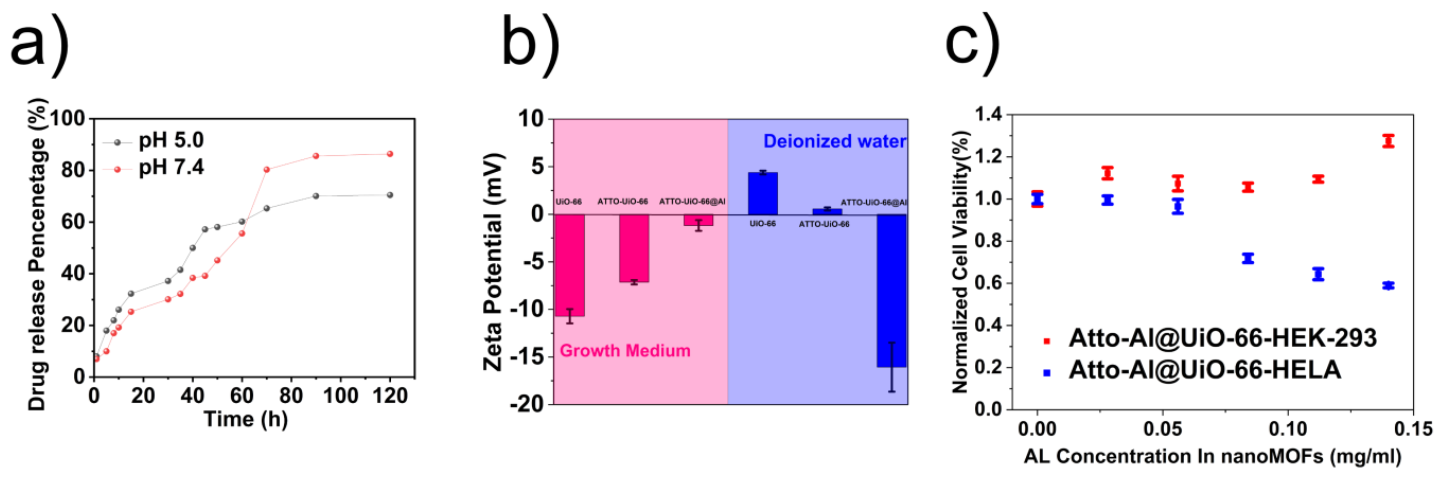
Assessment of defect-engineered nanoMOFs for cargo release and cytotoxicity: a) Quantification of the time-dependent release percentage of AL from ATTO-UiO-66@Al in pH = 5.0 (based on HCl) and pH = 7.4 PBS solution depending on the UV-Vis adsorption at 205 nm. b) The Zeta potential of UiO-66, ATTO-UiO-66, and ATTO-UiO-66@Al in cell growth medium (10 % FBS) or deionized water. c) MTT assay for cell viability (%) for different concentrations of AL loaded in ATTO-UiO-66, data recorded for 48 h after incubation in HeLa cells and HEK-293 cells (n = 4).

It is noteworthy that the released amount reached 85 % within 90 h at pH 7.4, while the corresponding value was less than 70 % within the same time duration at pH 5.0. Such a distinctive alteration in drug release behavior could be attributed to the lower degradation rate of UiO-66 based on HSAB (hard and soft acid and based) theory^[22]^ under acidic conditions than under neutral and basic conditions. The zeta potential results show there is an amount of drug loaded in the nanoMOFs and the surface will be coated by a protein corona and change the zeta potential of particles in cell growth medium compared within water, which offers a protective layer for the drug delivery (Figure 6b).

To test in vitro cytotoxicity of the UiO-66 NPs, cell viability was examined by standard MTT assays against HEK-293 and HeLa cells. Interestingly, free AL (Figure S24) appears cytotoxic for both HEK-293 and HeLa cells. Incubation of ATTO-UiO-66 NPs loaded with AL on HEK-293 cells resulted in minor or no variation of cell viability (Figure 6c). This phenomenon remained consistent regardless of the concentration of drug loading (Figure 6c), underscoring the favorable biocompatibility of ATTO-Al@UiO-66 nanoparticles in vitro. It is noteworthy that ATTO-UiO-66@Al exhibited no cytotoxic effects on HEK-293 cells at concentrations of up to 500 µg/mL based on nanoMOFs. In contrast, the incubation of Hela cells with ATTO-Al@UiO-66 resulted in a decrease in cell viability (Figure 6c and Figure S25). The increase IC_50_ (see MTT Assay methods and Figure S26) in the effect of Al upon delivery from ATTO-Al@UiO-66 (IC_50_ = 151± 18.3 μg/mL) corresponds to delivered concentration much higher than that of the free drug (IC_50_= 3.3 ± 0.22 μg/mL). The bis-phosphonate configuration of AL is likely to strongly adhere with the Zr sites within the MOFs. Consequently, incomplete release could mitigate cytotoxicity and offer the potential for gradual, controlled release to alleviate in vivo side effects.

## 3 Conclusion

This research highlights the significance of defect engineering in UiO-66 Metal-Organic Frameworks (MOFs) with controlled defect density to enhance fluorescence modification and facilitate their loading with pharmaceutics for delivery in cells. Density Functional Theory (DFT) calculations are employed to assess the chemical stability of ATTO 655 when bonded to the surface of UiO-66 MOFs, a critical factor for long-term tracking and colocalization capabilities of ATTO-UiO-66 with subcellular organelles. By leveraging spinning disc microscopy the stable coupling of ATTO 655 on the modification of defect sites in MOFs’ was shown, showcasing their functionality and safety for biomedical applications, particularly in drug carrier delivery and live cell imaging.

Cutting edge single-particle tracking (SPT)^[7,8,9,10]^ and advanced machine learning analysis toolboxes, provided the first direct observations of the spatiotemporal localization of nanoMOFs. Coupled with a new colocalization algorithm, the data elucidate crucial insights into nanoMOFs’ internalization in HeLa cells and their colocalization with endo-lysosomal pathways. This is pivotal for comprehending the intracellular interactions of MOFs in real-time and their trafficking pathways. The novel approach, which combines defect engineering with advanced analytical methods, represents significant progress in targeted drug delivery and cellular imaging safely.

## Supporting Information

Supporting Information is available from the Wiley Online Library or the author.

## Acknowledgments

This work was supported by the Villum Foundation Center BIONEC (18333), the NNF Challenge Center for Optimised Oligo Escape and Control of Disease (NNF23OC0081287), the NNF Center for 4D Cellular Dynamics (NNF22OC0075851), the Lundbeck foundation grant R346-2020-1759, Villum foundation Synergy grant (40578) and Villum experiment grant (40801), and Carlsberg foundation grant CF21-0499. N.S.H. is a member of the Integrative Structural Biology Cluster (ISBUC) at the University of Copenhagen and an associate member of the Novo Nordisk Foundation Center for Protein Research, which is supported financially by the Novo Nordisk Foundation (NNF14CC0001). Funding from the China Scholarship Council (No. 202009350012) for this work is also acknowledged.

## Data and Code availability statement

All code is written in Python, code and data can be available from the corresponding author upon reasonable request.

