## Supplementary materials and methods for "Defect-Engineered Metal-Organic Frameworks as Nanocarriers for Pharmacotherapy: Insights into Intracellular Dynamics at The Single Particle Level"

### 1. Materials and Instrumentation

All reagents and solvents used in this study were obtained from commercial sources and used as received without further purification. PXRD (Powder X-ray Diffraction) analysis was conducted using a Rikagu Miniflex 600 Benchtop X-ray diffraction instrument. TGA (Thermogravimetric Analysis) was performed on a Seiko S-II instrument, where dried crystalline samples were heated at a rate of 5 °C/min up to 1000 °C and then cooled to room temperature under an $N_2$ atmosphere. $N_2$ isotherms of the samples were measured using an ASAP 2020 and 2460 instrument from Micromeritics Co. Ltd. Scanning Electron Microscopy (SEM) was carried out using the FEI Quanta 3D FEG Scanning Electron Microscope. Inductively Coupled Plasma (ICP) spectroscopy was conducted on a JobinYvon Ultima2 instrument. X-ray Photoelectron Spectroscopy (XPS) measurements were performed using a Thermo ESCALAB 250 spectrometer with a non-monochromatic Al Kα X-ray excitation source. The data obtained were analyzed using CaSa XPS software with the C 1s line (284.8

eV) as the reference line to compensate for surface-charging effects. $^1$H-NMR spectra were recorded using Bruker AVANCE III 400 MHz and JNM-ECZ400S spectrometers. Absorption spectra were measured using a UV-Vis absorption spectrophotometer (SHIMMADZU, Japan). IR spectra were collected using VERTEX 70 series FT-IR spectrometers. Elemental analyses were performed on an Elementar Vario MICRO Elemental analyzer. The Zeta Potentials of nanoparticles were measured using a BI-200SM Analyzer. Static water contact angles were characterized using a JC200D instrument. Three-dimensional images of particles and live cells were captured using the SDCM uses an oil immersion 60x objective (Olympus) and a numerical aperture of 1.4 connected to a CMOS camera (photometric PRIME 95B) with an effective pixel size of 183 nm x 183 nm. Single crystals of UiO-66 with dye under a bright field (MOF) and 488 nm laser (dye) were obtained using a Confocal laser scanning microscope Ti-E&C2 (Nikon, Japan). MTT assays were analyzed using a Plate reader iD3.

### 2. Experimental Section

#### 2.1 Synthesis of UiO-66 with different defect content

$ZrCl_4$ (100 mg) and BDC (Benzene-1,4-dicarboxylic acid) (150 mg) in 15 mL of N, N-dimethylformamide (DMF) solution was ultrasonicated for 10 minutes and followed by the addition of 3 g Benzoic acid. The obtained solution was ultrasonicated for an additional 15 minutes and transferred to some vials (3 mL for each). The vials were heated at 120 °C, 150 °C, and 180 °C in the oven for 24 h. The obtained powder was further washed with DMF three times and then with acetone three times to obtain UiO-66 with different defect content suitable for powder X-ray diffraction.

#### 2.2 Synthesis of defective colloid UiO-66

$ZrCl_4$ (100 mg) and BDC (Benzene-1,4-dicarboxylic acid)(150 mg) in 15 mL of N, N-dimethylformamide (DMF) solution was ultrasonicated for 10 minutes. The obtained solution was transferred to some vials (3 mL for each). The vials were heated at 60 °C for 4 days. The obtained powder was further washed with DMF three times and then with acetone three times to obtain UiO-66 with different defect content suitable for powder X-ray diffraction.

#### 2.3 Synthesis of single crystal of UiO-66

According to the reference[1] In a 20 mL vial, 12 mg (0.037 mmol) zirconium oxychloride octahydrate was dissolved in 1 mL DEF. Separately, 5 mg (0.03 mmol) of terephthalic acid was dissolved in 1 mL DEF. The solutions were mixed and 2 mL of formic acid was added.

The resulting solution was shaken and placed in an oven at 408 K for 2 days. Block crystals were obtained in 69 % yield.

**2.4 Synthesis of Gal-single crystal UiO-66**

The Single crystal UiO-66 (5 mg) was suspended in 1 ml Gallocyanine (0.5 mg/ mL Deionized water) and sonicated for 10 minutes until well-dispersed. Then the obtained solution was heated for 24 hours at 50 °C. The obtained powder was further washed with DMF three times and then with deionized water three times to obtain Gal-Single crystal UiO-66.

**2.5 Synthesis of 120-150 °C/180 °C UiO-66, UiO-66-BDC-OH and UiO-66-BDC-Br**

120°C-UiO-66(100 mg) in 15 mL of N, N-dimethylformamide (DMF) solution was ultrasonicated for 10 minutes and followed by the addition of 150 mg BDC, BDC-OH or BDC-Br. Put them into a 150 °C oven for 24 h to get 120-150 °C UiO-66, UiO-66-BDC-OH and UiO-66-BDC-Br (150 mg BDC in 180 °C oven for 24 h and get 120-180 °C UiO-66 ). The obtained powder was further washed with DMF three times and then with acetone three times to obtain UiO-66 suitable for powder X-ray diffraction.

**2.6 Gas Sorption Measurements**

The as-synthesized samples were soaked in acetone for 2 days with the supernatant being replaced by fresh acetone about every 10 h during the process to exchange and remove nonvolatile solvates (DMF). After removal of acetone by centrifugation, the samples were activated under vacuum at room temperature for 6 h and then dried again by using the “outgas” function of instruments at 100 °C for 10 h prior to gas adsorption. $N_2$ isotherm measurements were performed at 77 K to the pressure of 1 bar.

**2.7 Synthesis of Gal-120/150/180°C -UiO-66/Nano-UiO-66**

The activated 120/150/180°C -UiO-66/Nano-UiO-66 (5 mg) was suspended in 5 ml Gallocyanine (0.5 mg/ mL Deionized water) and sonicated for 10 minutes until well-dispersed. Then the obtained solution was heated for 24 hours at 50 °C. The obtained powder was further washed with DMF three times and then with deionized water three times to obtain Gal-120/150/180°C -UiO-66/Nano-UiO-66.

**2.8 Synthesis of ATTO-UiO-66**

The activated nanoUiO-66 (10 mg) was suspended in 1μg ATTO-655 (2 mL PBS solution) and sonicated for 10 minutes until well-dispersed. Then the obtained solution was heated for 24 hours under 50 °C. The obtained powder was further washed with DMF three times and then with deionized water three times to obtain ATTO-UiO66.

**2.9 Synthesis of ATTO-UiO66@AL**

The activated nanoUiO-66 (10 mg) was suspended in 20 mL Alendronate (10 mg/mL pH 4.8 (HCl aqueous solution) and sonicated for 10 minutes until well-dispersed. Then the obtained solution was heated for 24 hours under 50 °C. The obtained powder was further washed with DMF three times and then with deionized water three times to obtain UiO66@AL. The powder collected was measured by ICP to determine the AL amount in nanoMOFs. In order to get ATTO-UiO-66@AL, The activated nanoUiO-66 (10 mg) was suspended in 20 mL Alendronate (10 mg/mL pH 4.8 (HCl aqueous solution) and sonicated for 10 minutes until well-dispersed. Then the obtained solution was heated for 24 hours under 50 °C. Keeping the mother liquor and adding ATTO 655(10 μg) into the previous solution and sonicated for 10 minutes until well-dispersed. Then the obtained solution was heated for 24 hours at 50 °C. The obtained powder was further washed with DMF three times and then with deionized water three times to obtain ATTO-UiO66@AL.

**3.0 Cell culture**

Culturing of Cells. HeLa cells and HEK-293 cells separately were cultured in Nunc EasYFlask 25 $cm^2$ cell culture flasks with Dulbecco's Modified Eagle's Medium with 4500 mg/L glucose supplemented with 100 IU/mL penicillin, 100 μg/mL streptomycin, 2 mM L-glutamine, 0.01 mM nonessential amino acids, 1 mM sodium pyruvate, and 10% (v/v) FBS at 37 °C and 5% $CO_2$. At a confluence of 80−90% in the culturing flasks, HeLa cells were detached by 3x trypsin-EDTA treatment (5 mg/mL trypsin and 2 mg/mL EDTA in DPBS, pH 7.4) and seeded for subsequent experiments. HeLa cells were seeded at a density of $2.5 \times 10^4$ cells/well in 96-well plates for cell viability assessment, 24 h before the experiments using passages 3 to 10. $2.5 \times 10^4$ cells/well in 8-well plates for an internalized experiment on the spinning disk microscopy, 24 h before the experiments using passages 3 to 10.

Cellular specimens are prepared for microscopic analysis during subculturing. Post-centrifugation, cells are enumerated using a hemocytometer and seeded in Ibidi IbiTreat 8-well plates at 25,000 cells per well (HeLa and HEK 293) and incubated at 37°C, 5% $CO_2$ for 24 hours in growth media using passages 15 to 20. Pre-imaging, labeled cells(CellMask™ Green

Plasma Membrane Stain, Early/Late Endosomes-GFP, and LysoTracker® Green DND-26) undergo triple washing with 10 mM HEPES in HBSS, followed by the addition of 200 µL growth media with 10 µg/ml ATTO-UiO-66. Endosome labeling involves adding 1 µL CellLight™ (Early/Late Endosomes-GFP, BacMam 2.0) per 10,000 cells and incubating for 16 hours. Membrane labeling uses CellMask™ Green Plasma Membrane Stain (1:200 ratio) for 10 minutes. For lysosome labeling, LysoTracker® Green DND-26 is diluted 1:20,000 and incubated for 3 hours. For SPT of ATTO-UiO-66 and compartments was performed using 30 % 640 nm laser power for ATTO-UiO-66 and 10 % 488 nm laser power for endo/lysosomal compartments with 30.04 ms exposure time.

**3.1 Effect of free drug and ATTO-UiO-66@AL on cell viability**

For the cytotoxicity studies of free AL and AL-UiO-66, HeLa and HEK-293 Cells were seeded in a 96-well plate at a density of $2.5 \times 10^4$ cells per well and were incubated in DMEM media containing 10% fetal calf serum for 24 h at 37 °C. Then free AL and ATTO- UiO-66@ AL were added to the medium and the cells were incubated at 37 °C for 48 h. The drug concentrations were set as between the range 0-200 µg $mL^{-1}$ on an AL basis. The nanoparticle concentrations were also varied according to the drug concentrations. Cell viability was determined by the standard MTT assay. After 48h, Then HeLa cells with drug cultured in 96-well plates were washed twice with 37 ℃ HBSS (pH 7.4). Subsequently, HeLa cells were incubated with an MTT solution containing 240 µg/mL MTT in HBSS (pH 7.4) for 4 h at 37 °C and 5% $CO_2$. Terminate the culture, carefully aspirate the culture medium from each well, add 150 µl DMSO, and shake at low speed for 10 min. Then Absorbance was measured at 550 nm using an iD3 plate reader. The HEK-293 followed the same way. The MTT formazan can be imaged in different concentration drug-treated 96 well plates after MTT assay(under 50% 532 nm laser with 50.04 ms exposure time). The $IC_{50}$ value for ATTO-UiO-66@AL was determined through linear fitting based on HeLa cell viability on AL concentrations of 56 µg/mL, 84 µg/mL, 112 µg/mL, and 140 µg/mL. the $IC_{50}$ value for free AL, obtained through linear fitting based on cell viability on concentrations of 0 µg/mL, 1 µg/mL, 2.5 µg/mL, 5 µg/mL, and 12.5 µg/mL.

**3.2 Drug Release**

AL Release to measure in vitro drug release, two batches of 50 mg ATTO-UiO-66@Al were suspended in 5 mL of PBS (5 mM) with pH of 5.0 and 7.4. The suspensions were placed into pretreated dialysis bags with a molecular weight cut off of 3000 Da and sealed with dialysis bag holders. The sealed dialysis bag was put into a beaker with 50 mL of PBS with the same pH conditions. The beaker was shaken at 100 rpm at 37 °C. At certain time intervals, 0.2 mL of the release medium was taken out. The volume of the dissolution media was maintained at 50 mL. The AL concentration was measured spectrophotometrically by spectrophotometry by measuring the absorbance of Al (28 wt% loading as blank (14 mg / 50 mL )) against the test solution at wavelength 205 nm. These measurements gave information on the quantity of released AL.

Drug release %= 1 - ($Abs_{blank}$-$Abs_{Tn}$)/$Abs_{blank}$. (Tn : The time point at drug release)

#### 3.3 Single particle localization and Single particle linking

Upon successful localization and intensity extraction based on homemade software, particle trajectories need to be constructed by linking particles across frames. One commonly employed method is the nearest-neighbor algorithm utilized in multiple SPT frameworks[2][3]. This involves determining the distance between every particle in a given frame and those in the subsequent frame within a defined search range, which is the maximum distance a particle can travel within a single frame. The particle in the subsequent frame that displays the shortest distance is then considered the most likely position for linking the particle in the current frame. This procedure is repeated for all particles in all frames to create complete trajectories. However, certain experimental conditions can hinder this linking process from working perfectly. For instance, a particle may disappear either temporarily or permanently in a frame, leading to the termination of the trajectory. This temporary or permanent disappearance can arise due to factors such as photo-bleaching or movement of the particle away from the focal plane. In cases of temporary disappearance, a memory parameter can be set to account for gaps in particle visibility within frames. This parameter is utilized to address the effects of photoblinking or a particle moving away from the confocal plane and then re-entering it.

#### 3.4 Density Functional Theory Calculation

All periodic Density functional theory (DFT) calculations were performed using CASTEP [4]. The interactions between ion cores and valence electrons were described with PAW potentials. The electron exchange and correlation function was described by the Perdew-Burke-Ernzerhof (PBE) functional of generalized gradient approximation (GGA) and ultrasoft pseudopotentials were employed. The cutoff energy was set to 630 eV in our calculation. Brillouin zone integration was sampled with the 1 × 1 × 1 MonkhorstPack mesh k-point. The electronic iterations convergence was converged within $10^{-6}$ eV and all of the atoms were allowed to relax until the residual forces per atom were lower than 0.01 eV Å$^{-1}$. Though the PBE function may underestimate the energy band gap, it is still reliable to illustrate the variations of the band gaps and the relative activities of different naanoMOFs. The binding energies (ΔEad) were calculated by using Eq (1), in which Ead/surf, Ead, and Esurf were the total energies of the optimized adsorbates/surface system, the adsorbates in the gas phase, and the surface, respectively [5].

ΔEad = Ead/surf – Ead – Esurf (1)

**3.5 Methods section**

**Colocalization analysis**

Using the x,y, and t coordinates from nanoMOF detections, the endo/lyso channel was examined for particle-like signal at the MOF location by considering SNR and S/B_d, a signal intensity ratio.

S: the donut center intensity

B_d: the average intensity in donut interior

$$\text{SNR: } \frac{S-Bd}{\sigma(Bd)}$$

$$\text{S/B\_d: } \frac{S}{Bd}$$

The S/B_d ratio probes if there exists a maximum relative to the tail of a potential point spread function while the SNR may function as a measure of whether that maximum should be assigned to a particle or naturally occurring aberrations. For most conditions, three biological replicates were available (see Table 5 for specifications). The endo/lyso markers were taken as particles and hence colocalized with MOFs (since MOFs defined their origin) if both S/B_d > 1.1 and SNR > 1.1. The fraction of MOFs which colocalize with endo/lyso marker is computed in a detection basis, such that MOFs that colocalize in only parts of the track duration are not necessarily ruled out. The reported percentages of colocalization were calculated from the mean and standard deviation in the three biological replicates.

The percentages of colocalization naturally depend on the thresholds on S/B_d and SNR, though the procedure is still capable of reporting on relative differences among conditions.

**NanoMOFs Single Particle Tracking (SPT)**

MOFs were detected using a Laplacian of Gaussian detector with a 2-12 pixel range to accommodate for the size variation in NanoMOFs. The minimum allowed SNR was 0.8, which is low, though small entries are largely removed due to a subsequent filter demanding trajectories to be at least of a 7-frame duration. The search range was 8 pixels and the memory 0 frames.

Note that the 7-frame filtering in trajectory duration is still applied even when colocalization is calculated in a detection basis.

**Supporting Figures and Tables**

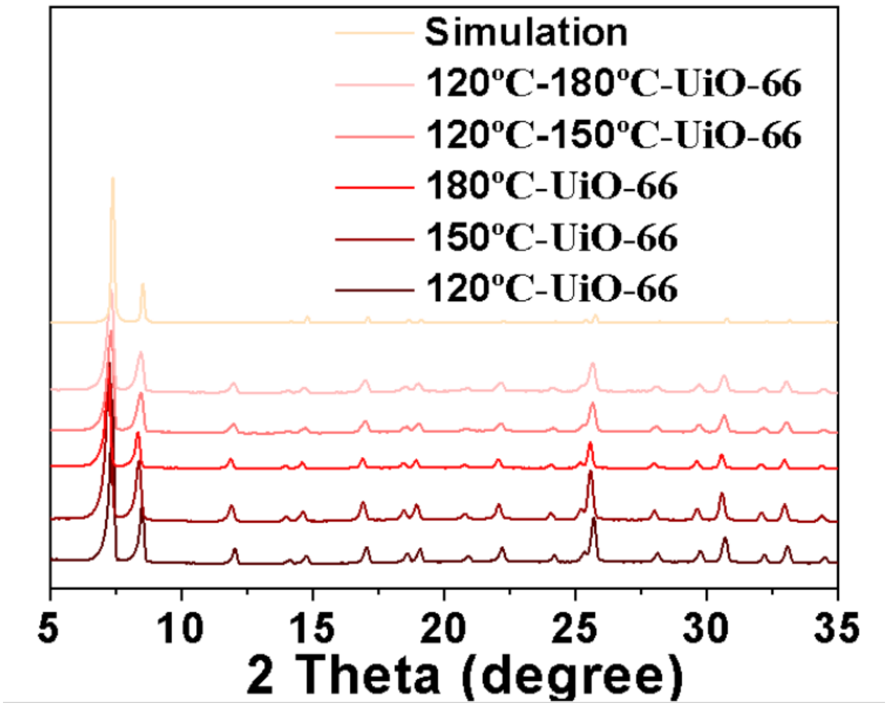


Figure S1 : PXRD of UiO-66 with different synthesis temperatures compared with simulation results.

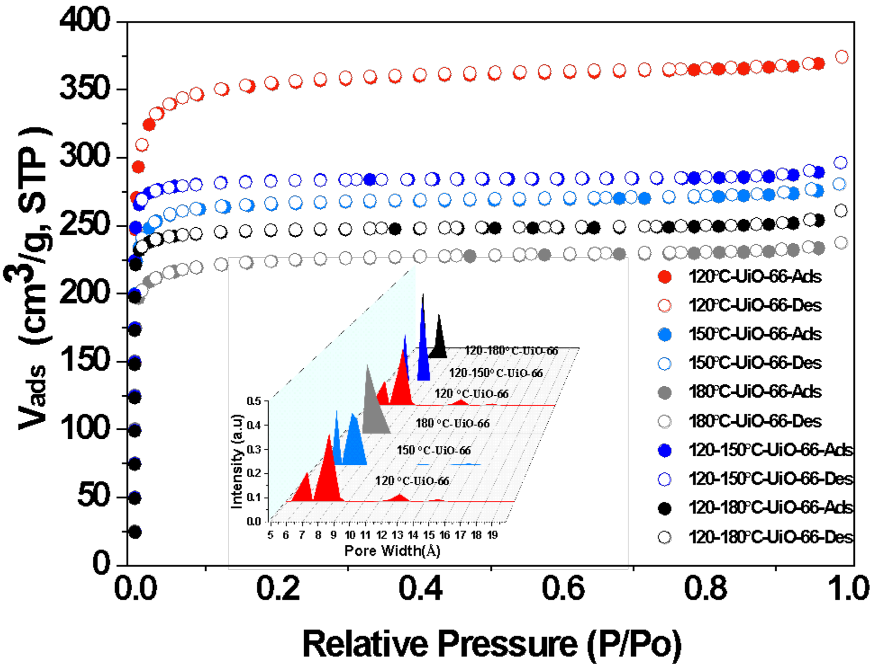


Figure S2 : $N_2$ uptake and pore size disruption of UiO-66 with different synthesis temperatures. .

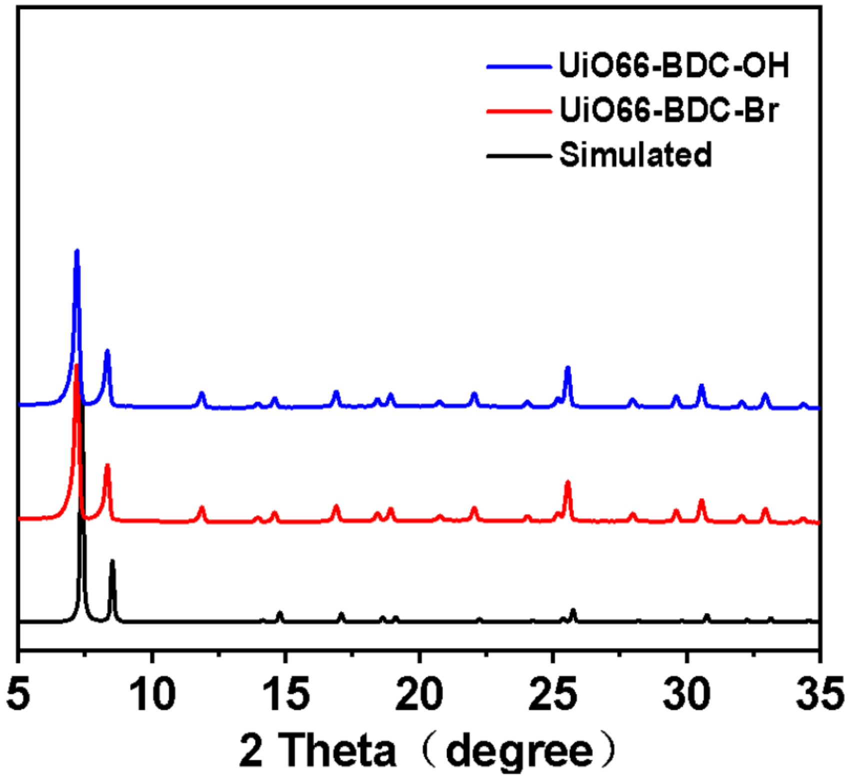

Figure S3 : PXRD of defective UiO-66 with BDC-OH and BDC-Br modification after post-synthesis method.

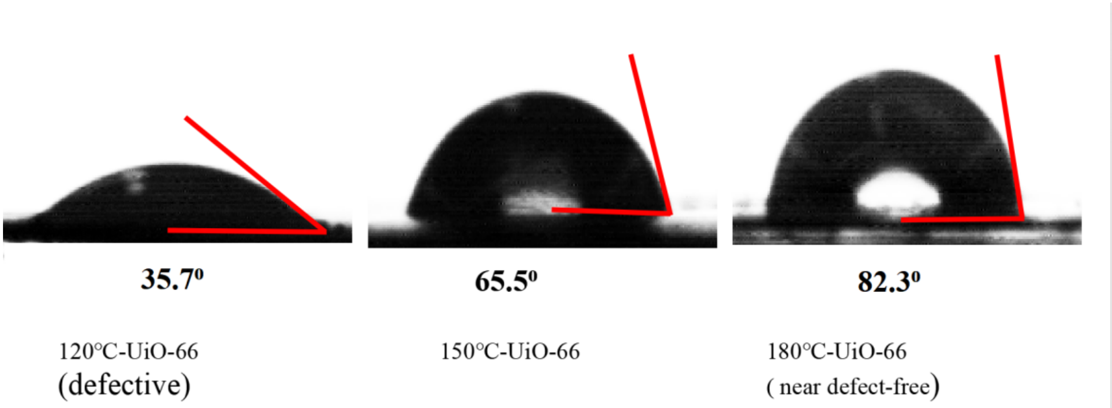


Figure S4 : The contact angle of different synthesis temperature MOFs.

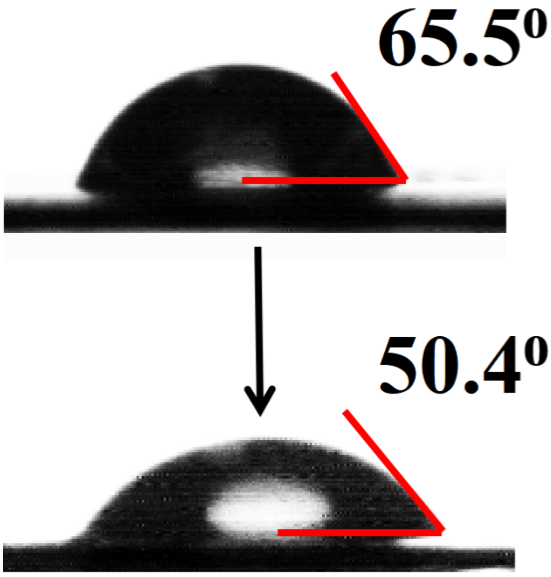


Figure S5: The contact angle of defective UiO-66 and BDC-OH modified defective UiO-66.

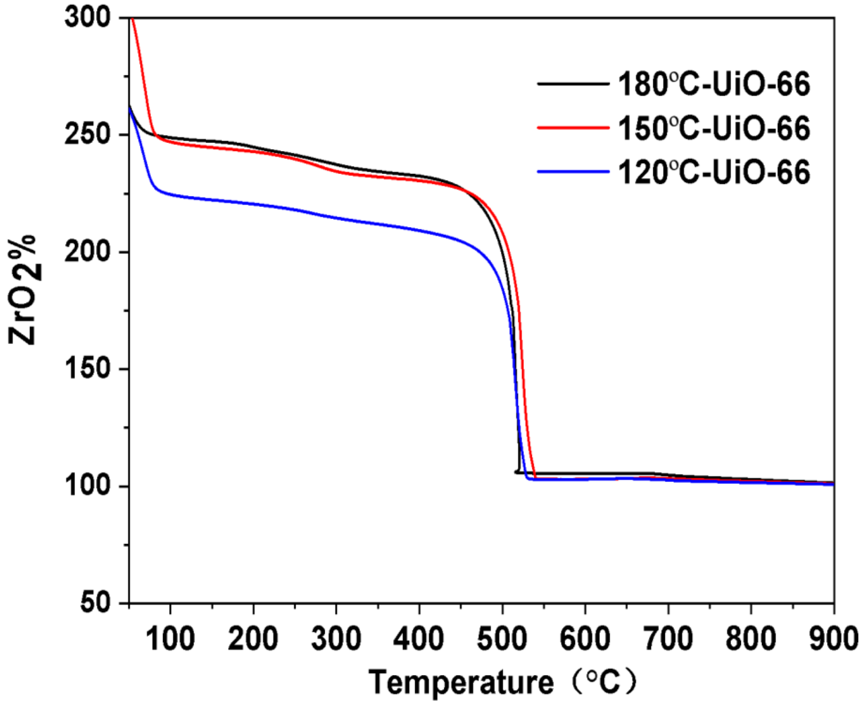


Figure S6: The TGA of different synthesis temperature MOFs.

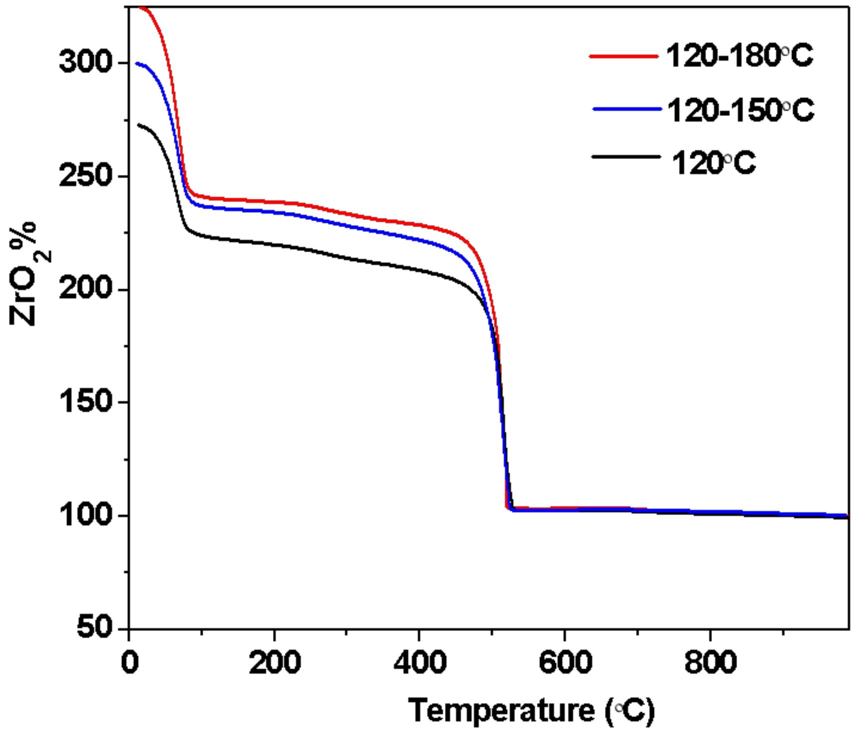


Figure S7: The TGA of different modified synthesis temperature MOFs.

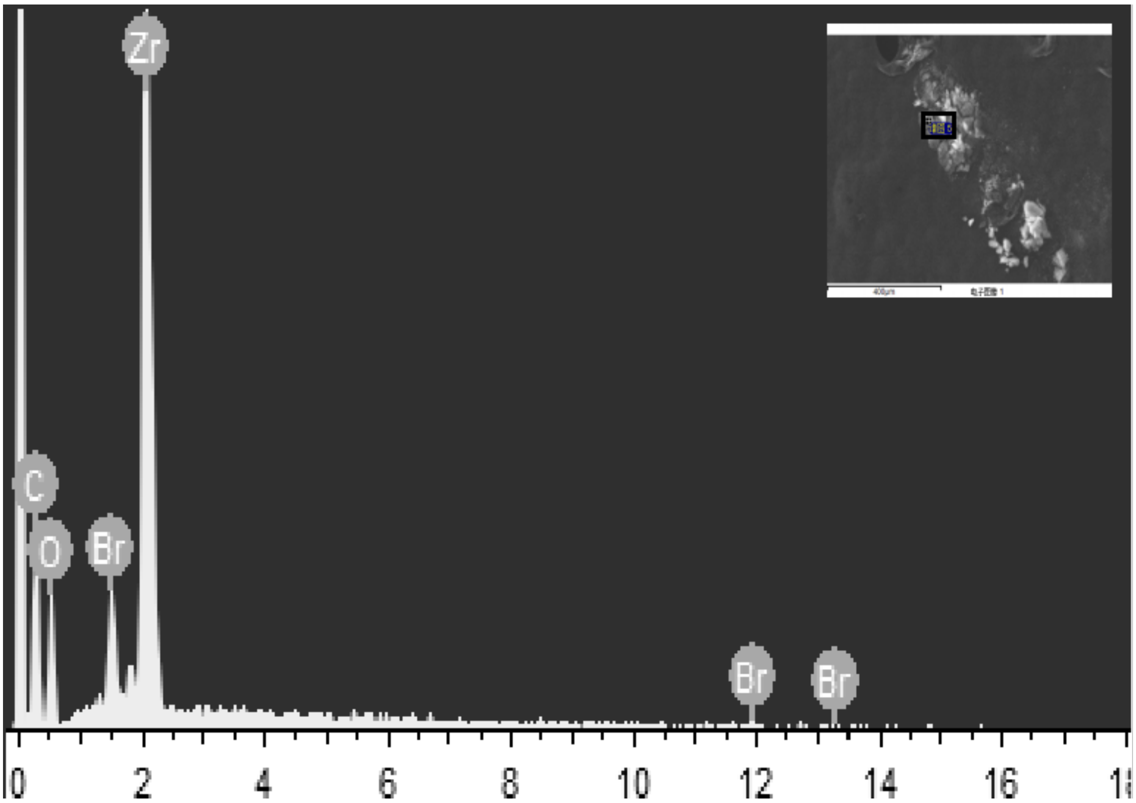

Figure S8: The XPS of defective MOFs modified by BDC-Br.

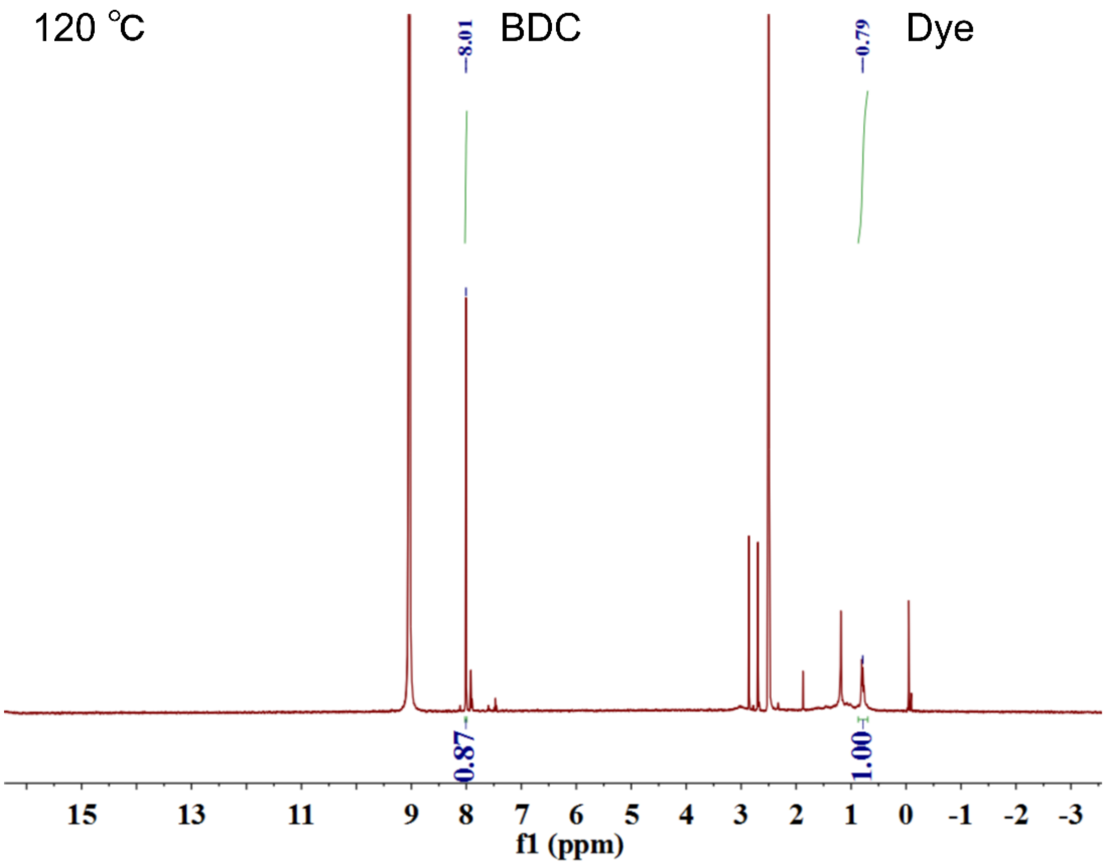
120 °C
BDC
Dye
8.01
0.79
0.87
1.00
15 13 11 9 8 7 6 5 4 3 2 1 0 -1 -2 -3
f1 (ppm)

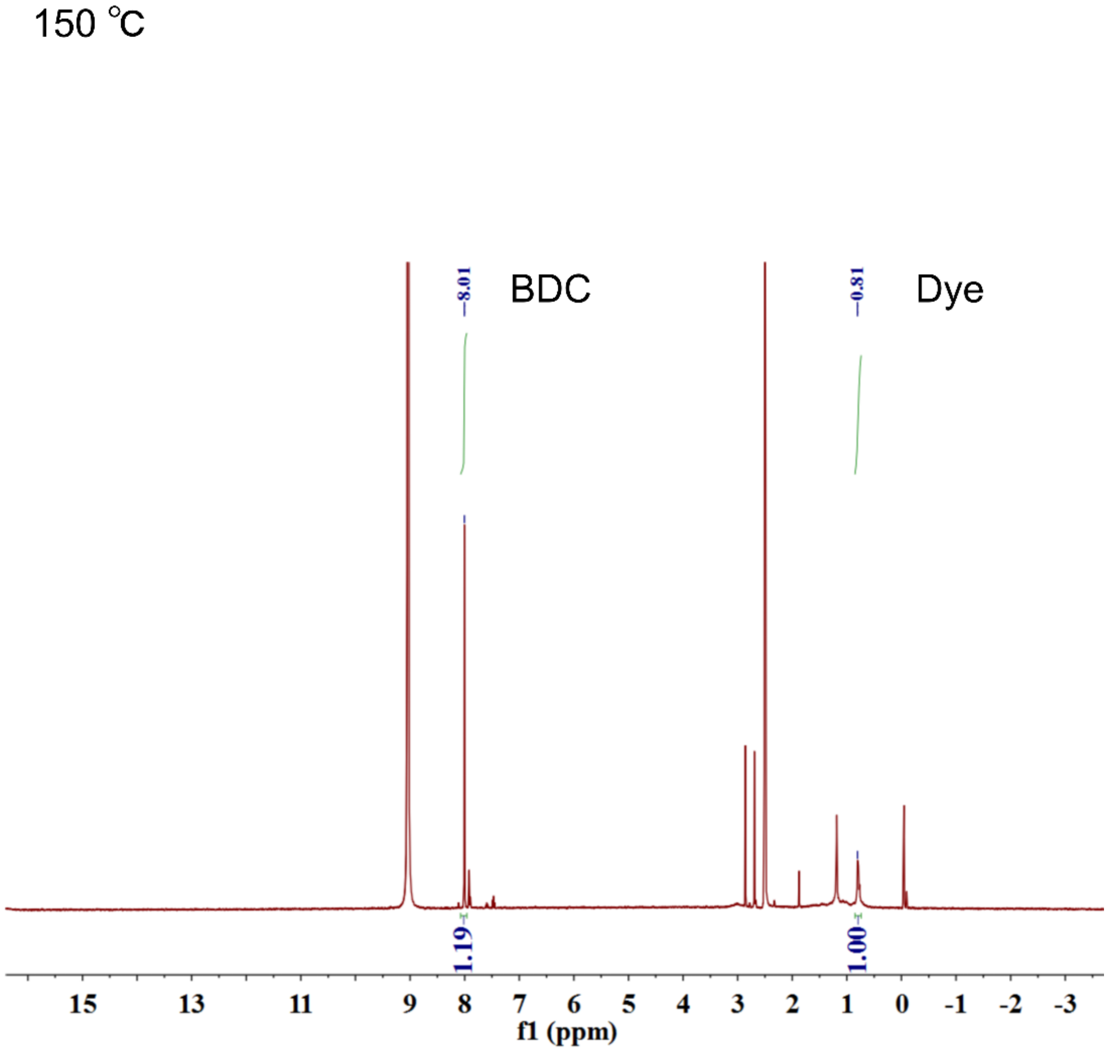
150 °C
BDC
Dye
8.01
0.81
1.19
1.00
15 13 11 9 8 7 6 5 4 3 2 1 0 -1 -2 -3
f1 (ppm)

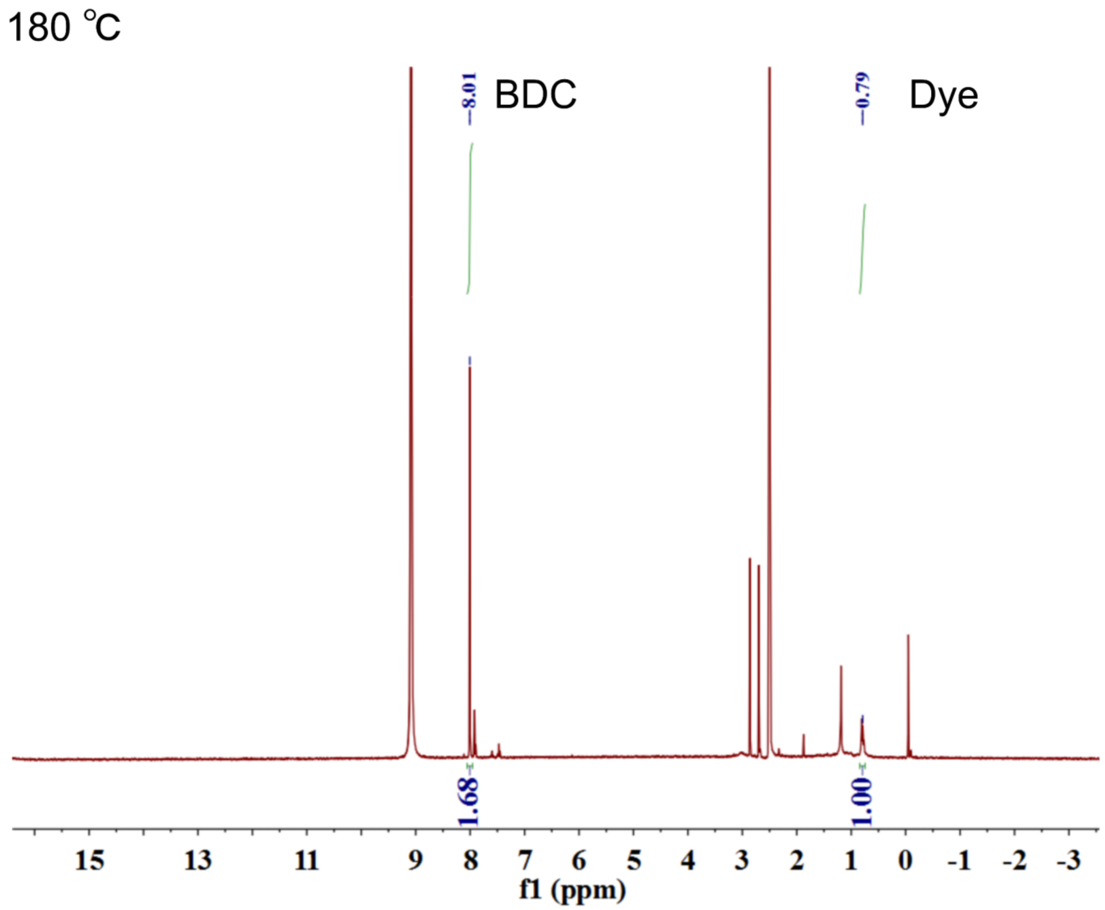


Figure S9: The [1]H-NMR of three types (different synthesis temperatures) defective MOFs modified by Gallocyanine.

| Temperature | 120°C | 150°C | 180°C |
|---|---|---|---|
| BDC (4H) | 0.87 | 1.19 | 1.68 |
| DYE (6H) | 1 | 1 | 1 |
| DYE:BDC（mol:mol) | O.766 | 0.560 | 0.397 |

Table 1: The DYE: MOF's ligand (mol: mol) of three types of defective MOFs modified by Gallocyanine based on [1]H-NMR results.

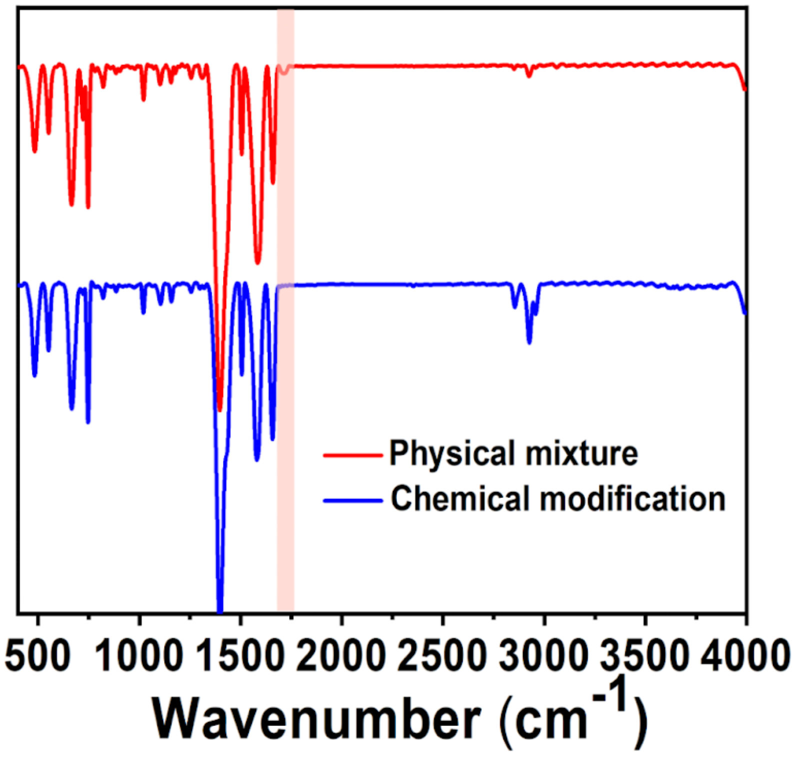


Figure S10: The FTIR spectra of physically mixed and chemically modified Gallocyanine with defective UiO-66. Red part marked (A rCOOH(C=O):Wavenumber=1750cm$^{-1}$)

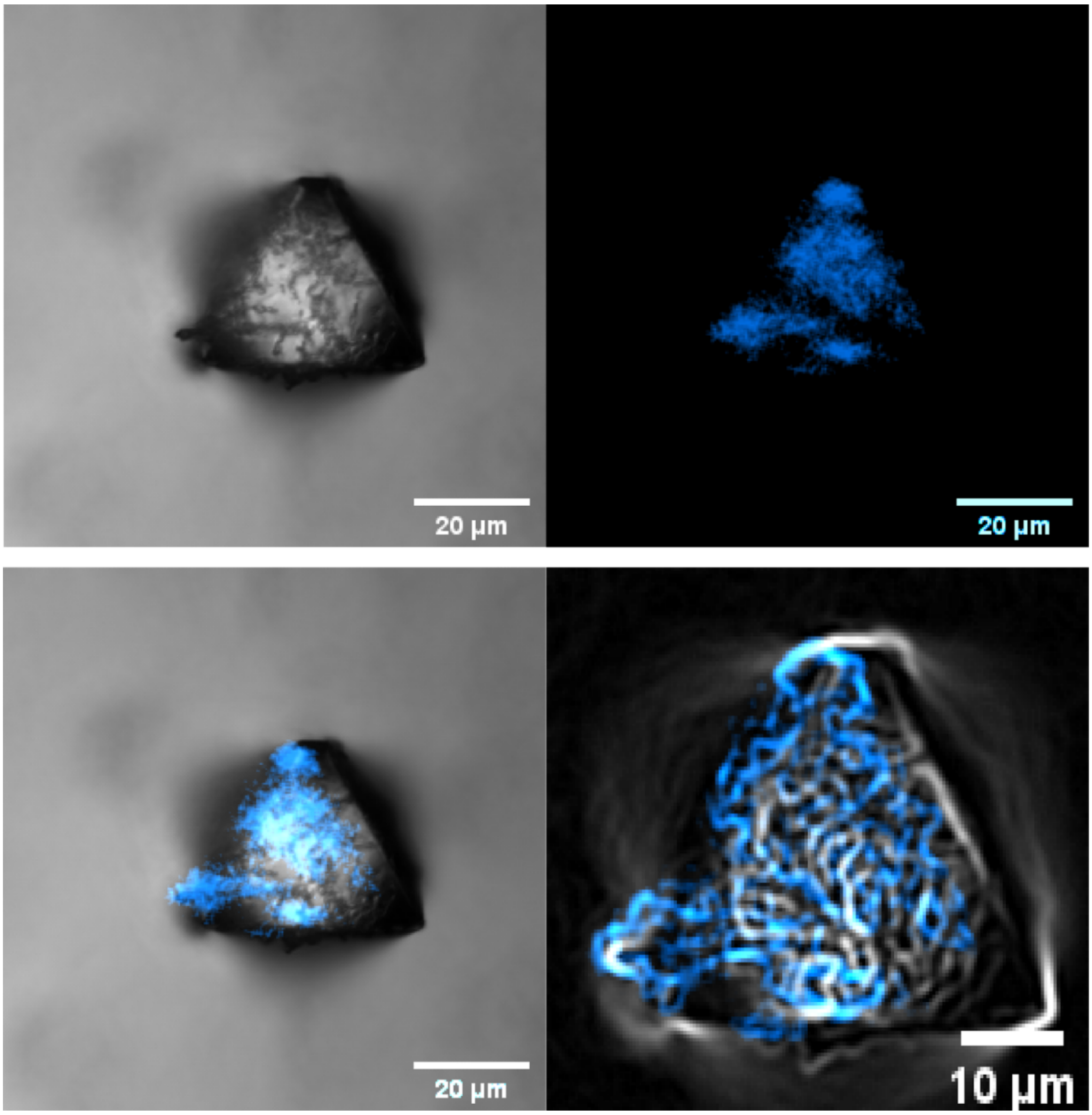


Figure S11: The single crystal images are arranged as follows: Top left: UiO-66 (bright), top right: modified by Gallocyanine (blue channel: 408 nm excited); bottom left: merged image showing both UiO-66 and Gallocyanine; bottom right: edge detection of the merged image.

| Temperature (°C) | BET ($cm^3$) | Contact Angel (°) | $ZrO_2$ % | Dye/BDC (mol: mol) |
|---|---|---|---|---|
| 120 | 1430.92 | 37.5 | 237 | 0.766 |
| 150 | 1068.8 | 65.5 | 234 | 0.56 |
| 180 | 895.06 | 82.3 | 214 | 0.397 |

Table 2: The results for the correlation plot between synthesis temperature (°C), BET ($cm^3/g$), contact angle (°), TGA (ZrO2 %), and the rate of dye/BDC (mol/mol).

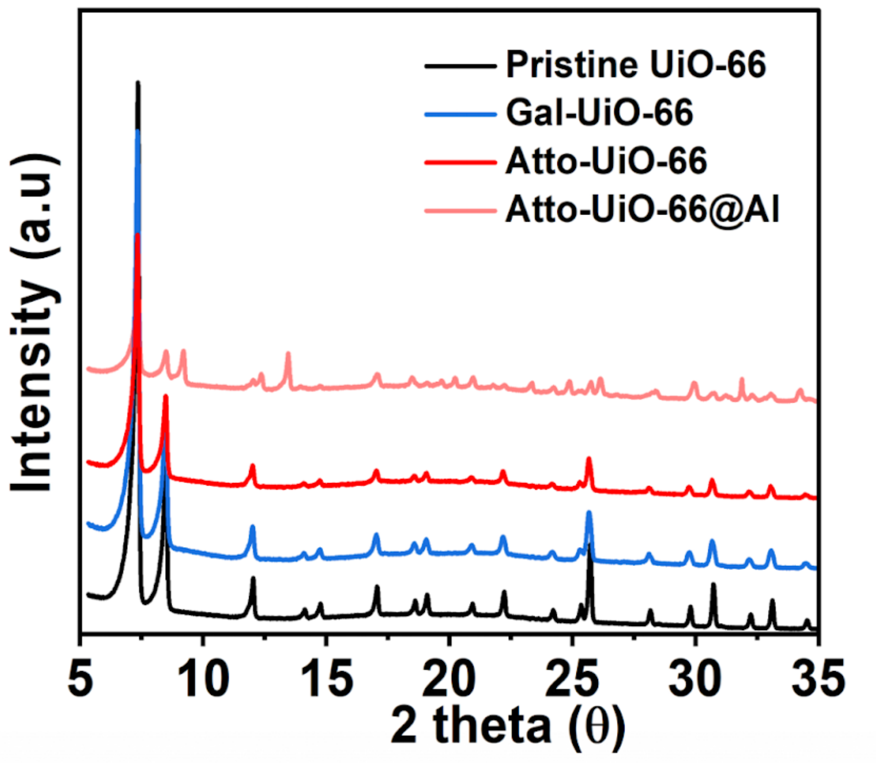


Figure S12: The PXRD of defective MOFs modified by ATTO 655, both ATTO 655 and Alendronate Sodium, and Gallocyanine.

**2.2 Density functional theory calculations displaying the ATTO 655 doping**

Depending on the missing linker, In the case of defective UiO-66, it is common to observe the absence of one ligand (11 SBU) or the absence of two ligands (10 SBU). For Structure I (Str I) (Figure S13a top), the defect structure with a missing linker was created by removing $BDC^{2-}$ linkers from the pristine UiO-66, and the charge was compensated by adding ATTO 655 molecules. Structure II (Str II) (Figure S13a bottom), the defect structure with two missing linkers were created by removing two $BDC^{2-}$ linkers from the pristine UiO-66. Para-doping was found to be the most stable, optimal existence when testing different Zr node positions modified by ATTO 655, whereas ortho-doping has significant steric hindrance and cannot set up the model. The distances between Zr and O (from ATTO 655) are as short as 2.229 Å and 2.206 Å, 2.220 Å for Str I and Str II, respectively, indicating the presence of chemical bonds between ATTO 655 and UiO-66. To understand the origin of charge in the electronic and optical properties of ATTO 655 and UiO-66, the band structures of pristine and ATTO 655-

doped one ATTO 655 and two ATTO 655 MOFs were calculated via the DFT method. Firstly, we optimized the structures of pristine UiO-66 with no doped ATTO 655 molecule, and the corresponding density of states (DOS) was determined (Figure S13b). For pristine UiO-66, the energy was predominantly localized on the linkers. We then obtained energy band gaps of 0.297 eV, 0.058 eV, and 0.031 eV for the pristine UiO-66, one ATTO 655-doped UiO-66, and two ATTO 655-doped UiO-66, respectively. For Str I, DOS is largely localized on ATTO 655, and the energy levels of ATTO 655 are introduced to the original DOS of pristine UiO-66, which decreases the DOS gap of the BDC Linker (Figure S13c). Similarly, Str II was optimized, and the calculated DOS (Figure S13d) showed a smaller band gap. These calculated results indicate that Str I and Str II would have a wider light absorption range by red shift compared with pristine UiO-66, and Str II has higher photogenerated electron-hole pair separation efficiency than pristine UiO-66 and Str I. As shown in Fig. S13e and S13f, there is a large overlap between the p/d orbitals of the Zr node and the p orbitals of oxygen form ATTO 655 in the region of -7–0 eV, again indicating that the dopant molecules were strongly bound at the Zr node. In addition, for Str I and Str II, the defect structure with a missing linker was created by removing one or two $BDC^{2-}$ linkers from the pristine UiO-66, and the charge was compensated by adding ATTO 655 molecules. The defect energy, i.e., the energy cost per removed linker, can then be calculated using Eq. (1). If the charge was compensated by adding phosphate molecules and drug AL, it can then be calculated using Eq. (2) and Eq. (3).

Eqn (1): E = E(Host : ATTO 655) - [E(Host) - E(Linker) + E(ATTO 655)]

Eqn (2): E = E(Host : PO43-) - [E(Host) - E(Linker) + E(PO43-)]

Eqn (3): E = E(Host : alendronate) - [E(Host) - E(Linker) + E(alendronate)]

The results show that ATTO 655 is more competitive in bonding to the Zr node on the surface of UiO-66 compared with phosphate (Figure S13g and Table 3) and AL (Table 3), which provides us with a feasible method to load the drug in the volume first and then modify the dye on the surface.

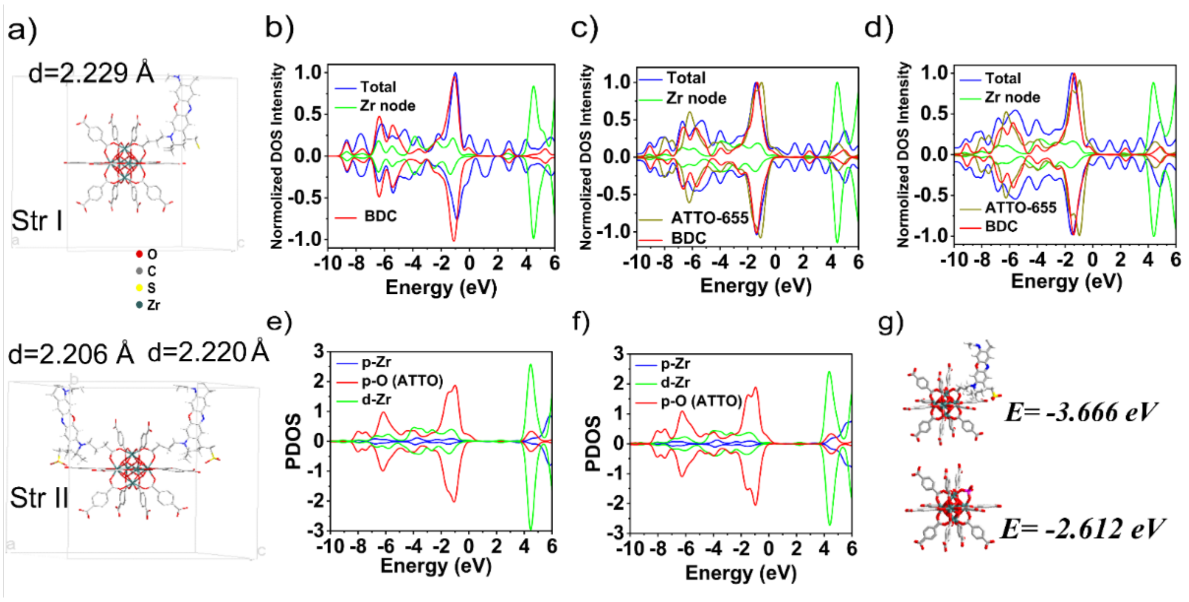


Figure S13: Density functional theory calculations displaying the ATTO 655 doping. a) Optimized structures Top: Str I and Bottom: Str II. The distances between node-Zr and ATTO-O is illustrated; Calculated density of states for the b) pristine UiO-66. c) ATTO-UiO-66 built as Str I. d) ATTO-UiO-66 built as Str II. e) Node-Zr and O-ATTO in Str I and f) Node-Zr and O- in ATTO Str II. g) The adsorption energy: ATTO 655 or phosphate with UiO-66´s Zr node.

| | eV | kj/mol | | eV | kj/mol | | eV | kj/mol |
|---|---|---|---|---|---|---|---|---|
| PBS (Phosphate) | -1966.08 | -1.89 | ATTO 655 | -8519.32 | -8.18 | Alendronate | -4603.04 | -4.42 |
| MOF | -44300.90 | -4.25 | MOF | -44300.77 | -4.25 | MOF | -44300.85 | -4.25 |
| MOF+PBS (Phosphate) | -46269.59 | -4.44 | MOF+ATTO | -52823.76 | -5.07 | MOF+Al | -48906.92 | -4.42 |
| Adsorption energy | -2.61 | -2.51 | Adsorption energy | -3.67 | -3.52 | Adsorption energy | -3.04 | -2.91 |

Table 3: The adsorption energy MOFs bonded with Phosphate, ATTO 655, and Alendronate based on DFT calculation.

| | | | | |
|---|---|---|---|---|
| | Inside | 6 | 0 | 44 |
| 4h | hetero+inside | 17 | 23 | 12 |
| | Outside | 35 | 5 | 80 |
| | Inside | 55 | 0 | 208 |
| 8h | hetero+inside | 175 | 39 | 28 |
| | Outside | 122 | 6 | 157 |
| | Inside | 71 | 14 | 282 |
| 24h | hetero+inside | 229 | 127 | 183 |
| | Outside | 201 | 130 | 41 |
| Biological replicates(BR) | | BR1 | BR2 | BR3 |

Table 4: The machine learning analysis of nanoMOF particle counts based on cell and particle trajectories for evaluation of nanoMOF trajectory quantities inside cells, hetero+ inside cells, and outside cells across three biological replicates in each condition is summarized in the table.

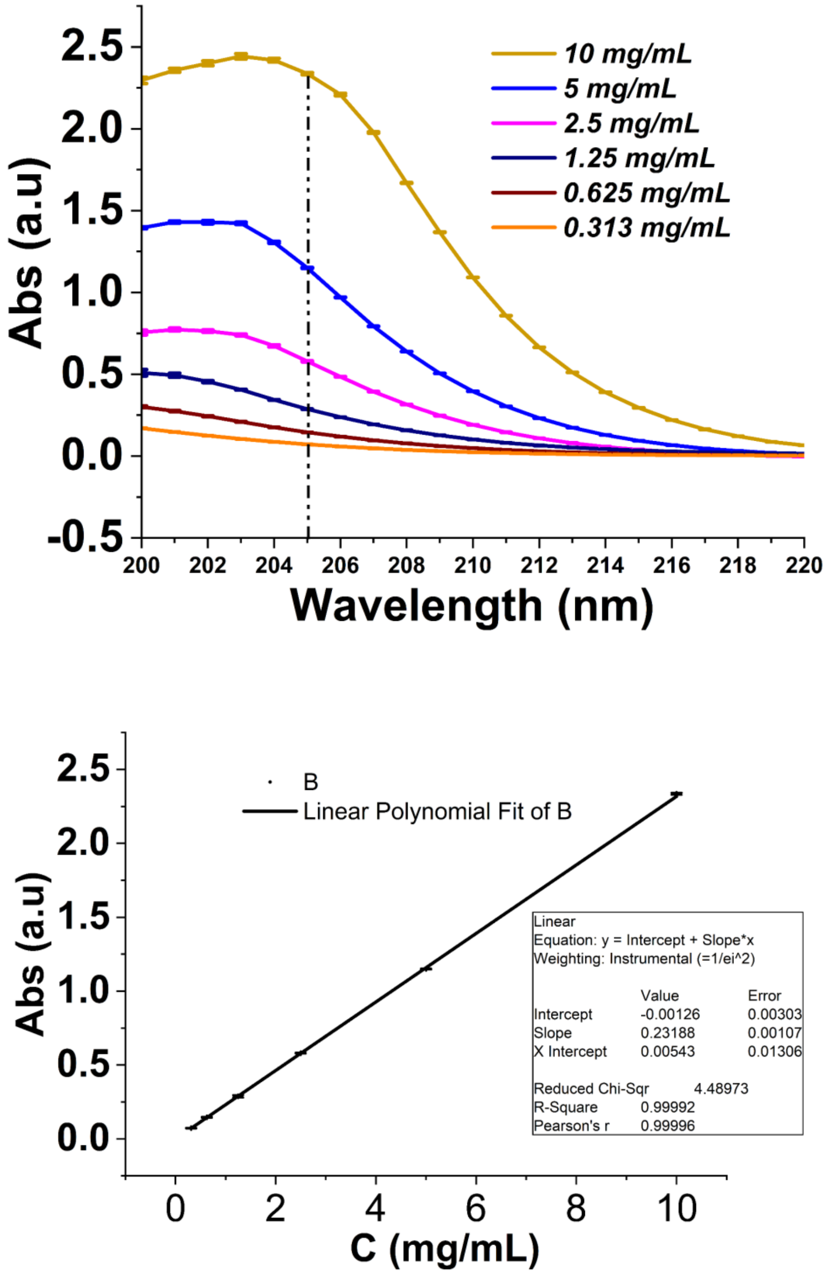


Figure S14: The intensity of Alendronate (AL) at different concentrations and the linear relationship between signal intensity and the concentration of Alendronate.

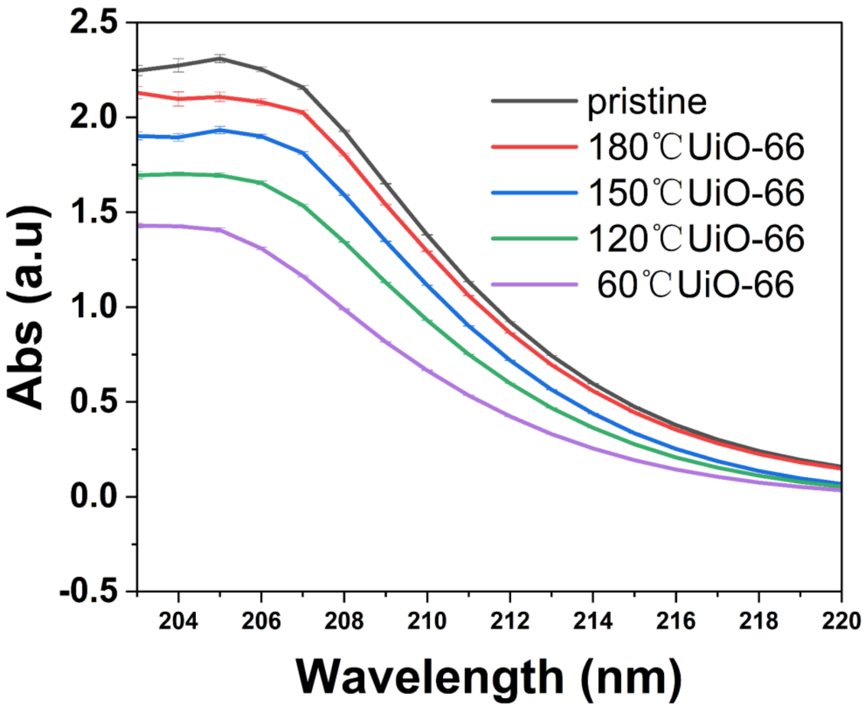


Figure S15: The Abs intensity of the Alendronate in upper water centrifuged after the different defective levels of UiO-66(10 mg) soaked in 20 mL (10 mg/mL pH 4.8 HCl aqueous solution) Alendronate for 24 hours at 50 ℃ (Wavelength = 205 nm). Drug Load %= ($Abs_{0h}$ - $Abs_{24h}$)/$Abs_{0h}$.

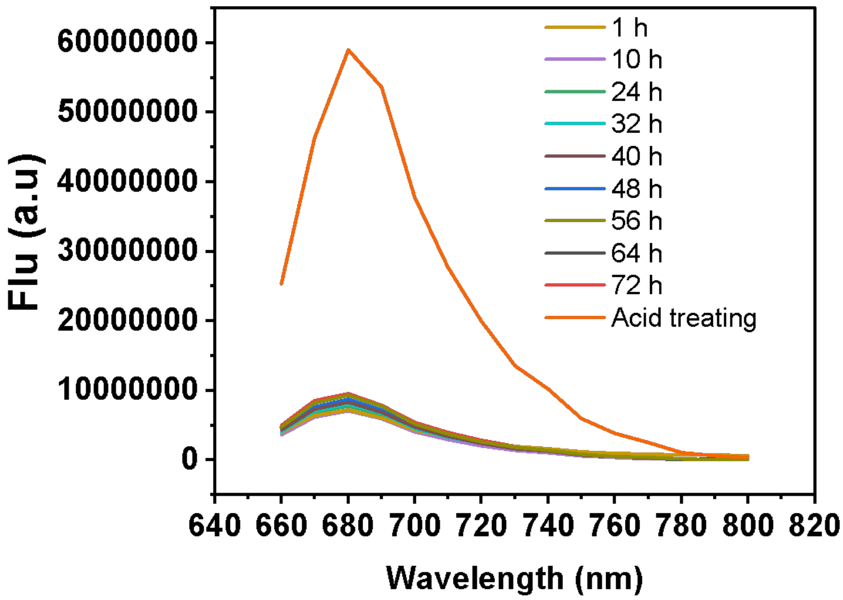


Figure S16: Soaking 5 mg ATTO-UiO-66 into 1 mL PBS solution, the signal intensity of ATTO 655 in the upper water layer monitor after centrifugation for 72 hours (excited at 600 nm).

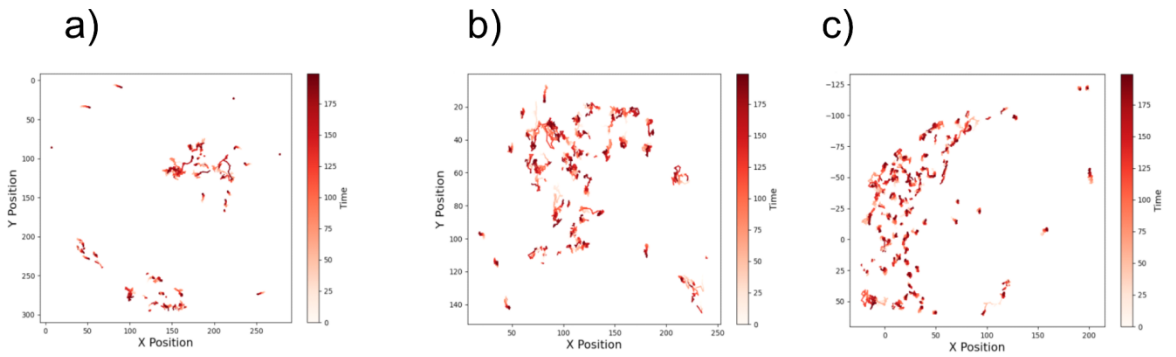


Figure S17: The internalized ATTO-UiO-66 trajectories in HeLa cells a) the ATTO-UiO-66 internalized in HeLa cells with a)early endosomes labeled with an expression of protein GFP Rab 5; b) late endosomes labeled with an expression of GFP Rab 7; c) lysosomes labeled LysoTracker® Green. (X,Y position unit = pix, 1 pix = 183 nm)(Time unit = Second).

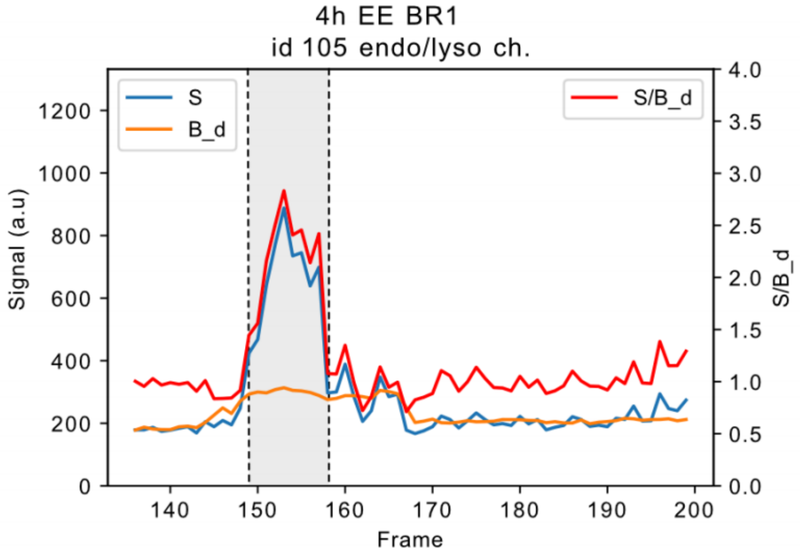


Figure S18: Time series of endo/lyso signals in nanoMOF particle center. S, the average local background endo/lyso signal from an encompassing donut, B_d, and their ratio S/B_d. S/B_d ratios of around one suggest that there is no endo/lyso signal colocalized with nanoMOFs. The displayed series is from 4h EE BR1 particle id 105, which exists in frame 132-200. At around frame 150, the interior signal, S, doubles while the background, B_d, is only slightly elevated. Considering the donut geometry, this is a period where MOFs colocalize with particle-like (radially decreasing from a center) signal.

a)

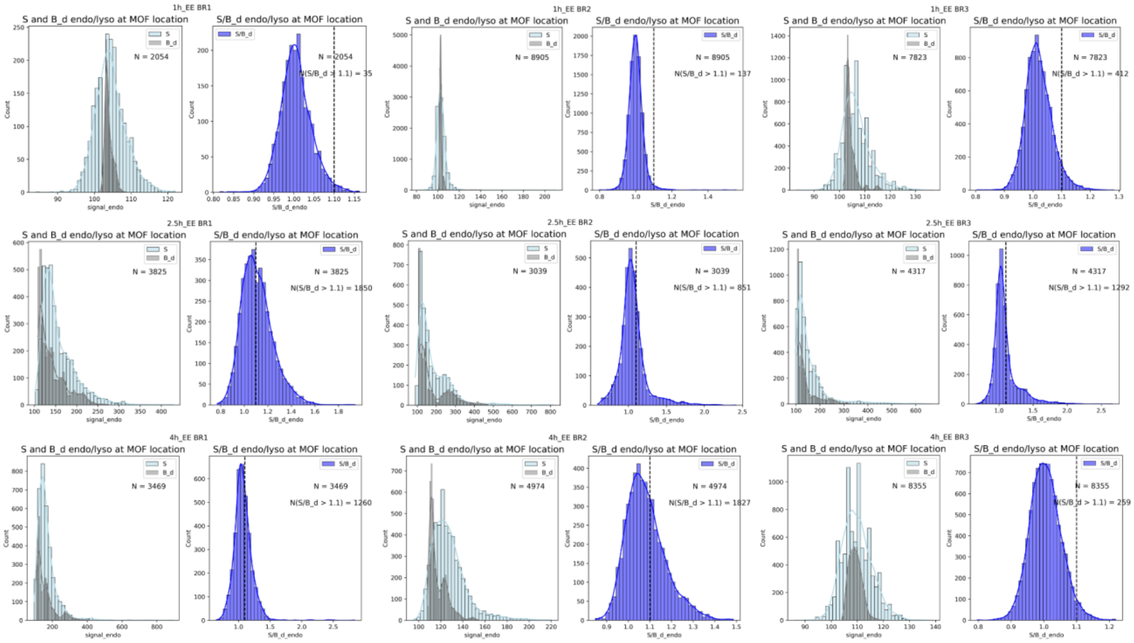


Figure S19: The counts of signal in S and B-d endo/lyso at MOF location and S/B endo/lyso at MOF location at 1h, 2.5h, and 4h with early endosome(3BR).

b)

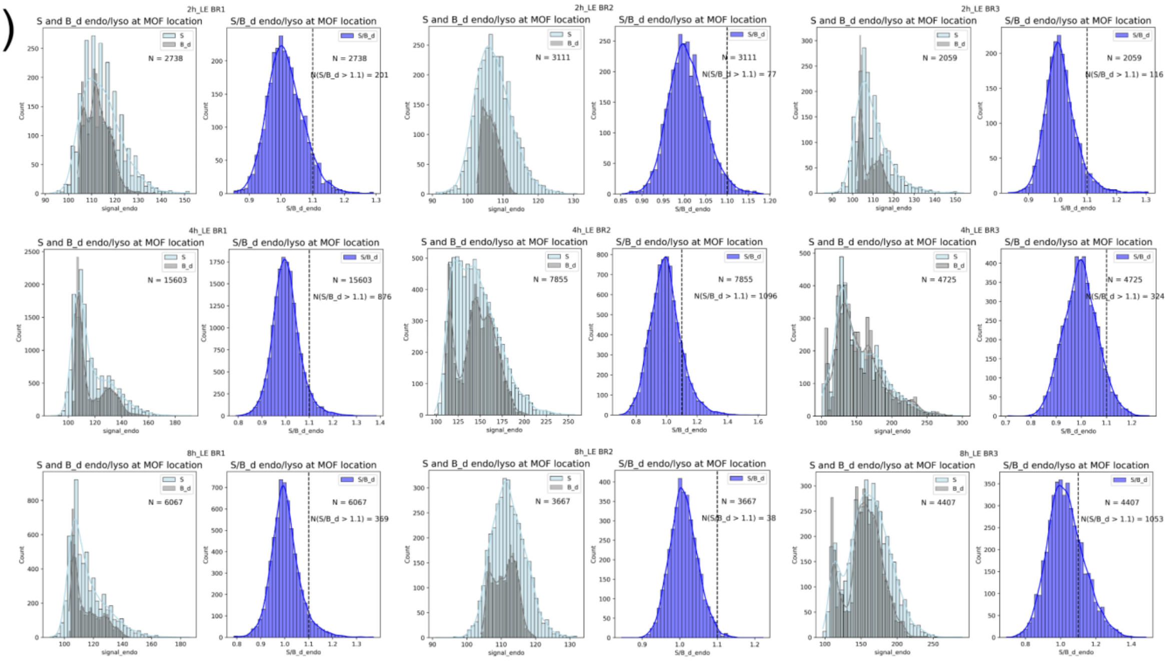


Figure S20: The counts of signal in S and B-d endo/lyso at MOF location and S/B endo/lyso at MOF location at 2h, 4h, 8h with Late endosome(3BR).

c)

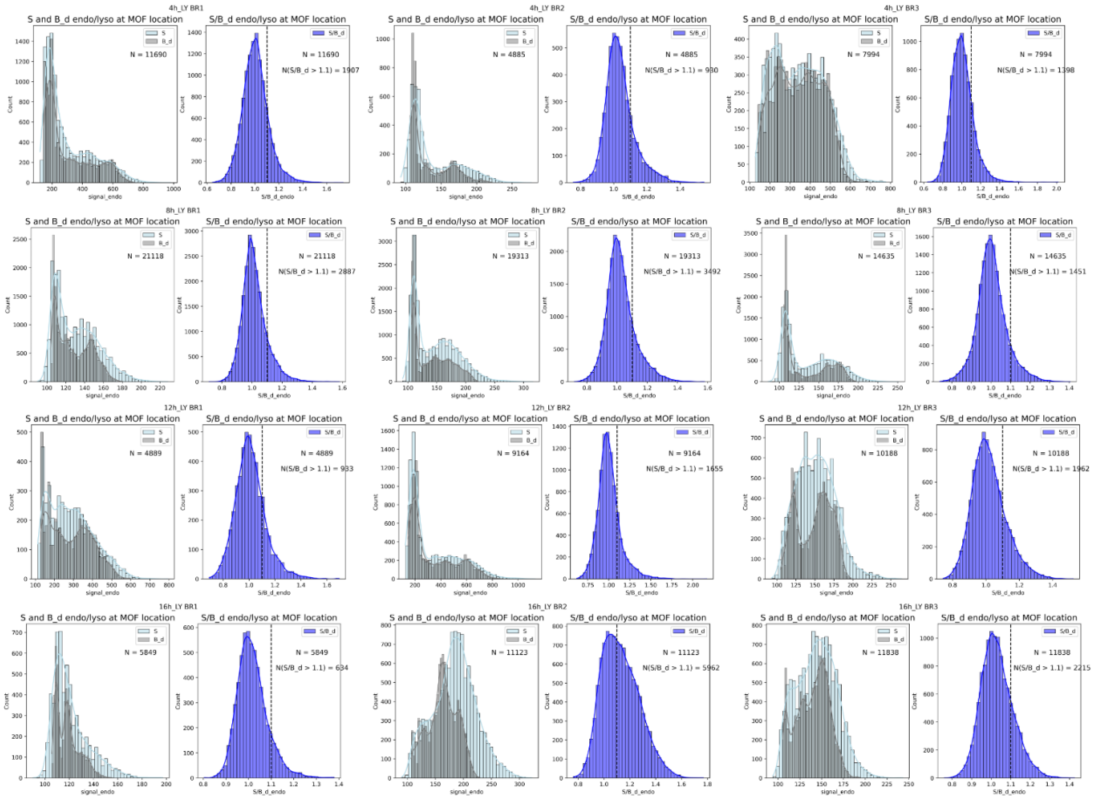

Figure S21: The counts of signal in S and B-d endo/lyso at MOF location and S/B endo/lyso at MOF location  at 4, 8h, 12h, 16h with Lysosome(3BR)

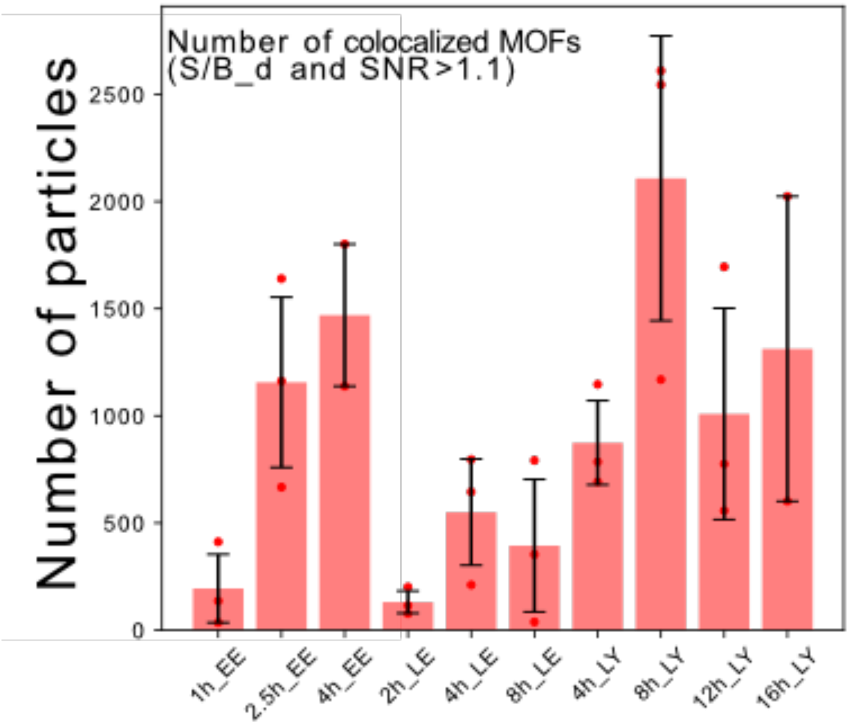


Figure S22: The number of colocalized nanoMOFs in different conditions.

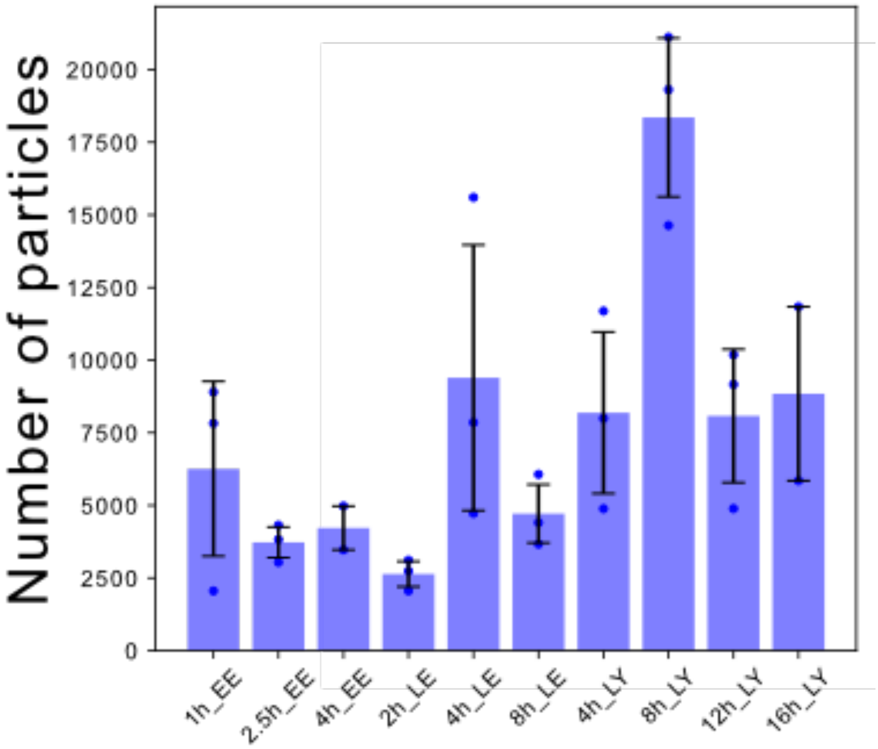


Figure S23: The total number of detection in FOV in different conditions.

The following attributes, d+, d-, s, etc. are qualitative and taken relative to a generalized imaging condition.

M: nanoMOFs channel

e/l: endo/lysosomal compartments channel

d+: increased particle density (can link particles location within an acceptable margin of error)

d-: decreased particle densities: sharp/higher SNR (can link particles location within an acceptable margin of error)

b: blurry/lower SNR

a: aggregated

os: oversaturated

* BR3 (4h EE) is discarded as the SNR in the e/l channel is lowered compared to BR1 and BR2 in the same condition.

** BR2 (16h LY) is discarded due to lower SNR in the nanoMOFs channel, maybe due to long time in lysosome causing linking bigger errors in tracking.

| Condition/BR | BR1 | | BR2 | | BR3 | | Assessment |
|---|---|---|---|---|---|---|---|
| | M | e/l | M | e/l | M | e/l | |
| 1h EE | d- | | | | | | acceptable |
| 2.5h EE | d- | s | d- | s | | s | good |
| 4h EE | a d+ | | a | b | | b | acceptable* |
| 2h LE | d- | b | d- | b | d- | b | acceptable |
| 4h LE | d+ | b | d- | b | | | acceptable |
| 8h LE | | b | a | b | d- | b | acceptable |
| 4h LY | d+ | b | | | d+ | b | good |
| 8h LY | d+ | b | d+ | b | d+ | b | good |
| 12h LY | | b os | | | | | good |
| 16h LY | | b | **b** d+ | b | | | good** |

Table 5: Assessment of microscopy data quality among the three biological replicates in each condition.

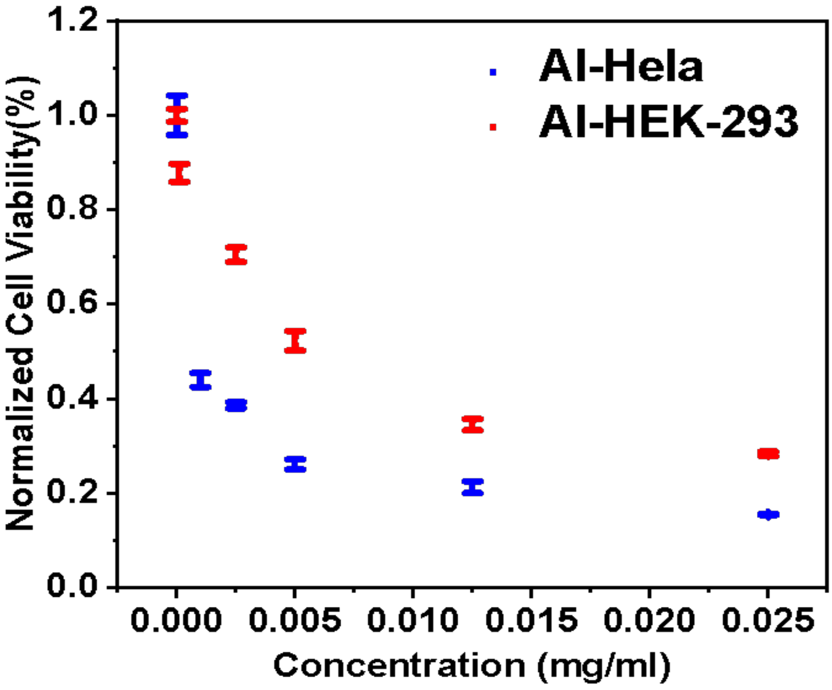


Figure S24: Normalized cell viability (%) for different concentrations of free Al in HEK-293 and HeLa cells after 48 hours of incubation (n=4).

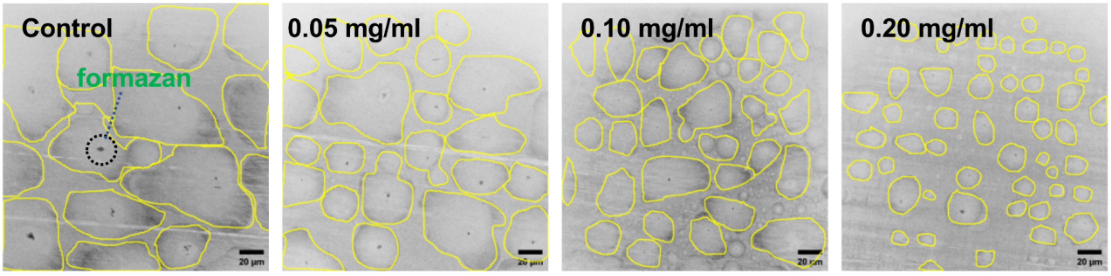


Figure S25: The 532 nm laser (excited MTT formazan's fluorescence) HeLa cells image control and with different drug concentrations in nanoMOFs (0; 0.05; 0.1; 0.2 mg/mL) 48 hours incubation after 4 hours' MTT assay (240 µg/mL) (Scale bar = 20 nm). Cell segmentation using cellpose.

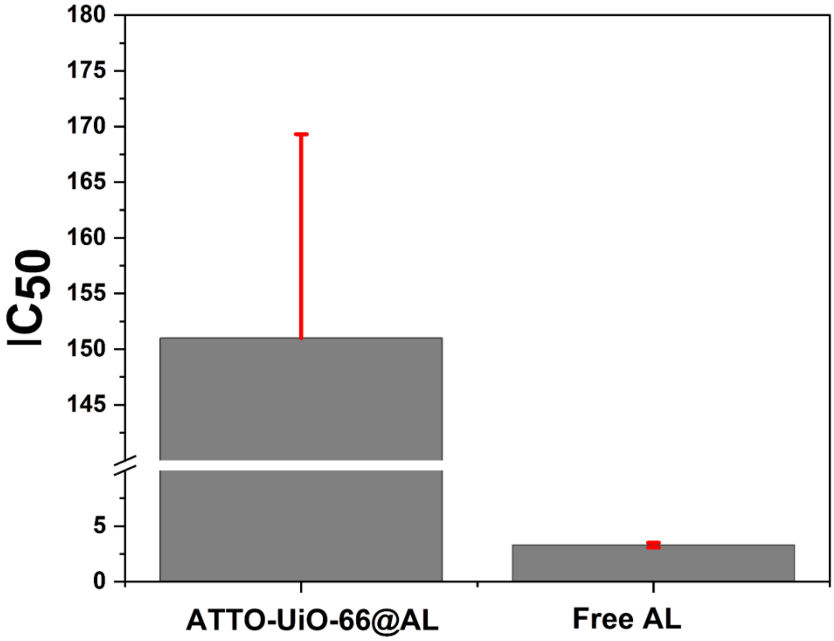


Figure S26: Comparison of $IC_{50}$ values of ATTO-UiO-66@AL (151.0 ± 18.3 µg/mL) and Free AL(3.3 ± 0.22 µg/mL).

Acknowledgments

We would like to express our gratitude to the research group led by Tianfu Liu from FJIRSM Cas for providing the nitrogen adsorption equipment and TGA equipment, and to the research group led by Xinxin Xiao from DTU for providing the PXRD and UV-Vis absorption spectrophotometer. Rui Peng from the Pharma department in KU for providing Zeta potential. Yuzi Zhao offered software to help with the DFT calculation. We acknowledge the Core Facility for Integrated Microscopy, Faculty of Health and Medical Sciences, University of Copenhagen for providing the SEM equipment.

[3] Daniel B. Allan, Thomas Caswell, Nathan C. Keim, and Casper M. van der Wel. trackpy:

Trackpy v0.4.1. 4 2018.

[4] S.J. Clark, M.D. Segall, C.J. Pickard, P.J. Hasnip, M.J. Probert, K. Refson, M.C. Payne, First principles methods using CASTEP, Zeitschrift Fur Kristallographie 220(5-6) (2005) 567-570.

[5] H. X. Liu, T. T. Liu, T. Huang, Z. B. Fang, L. Li, Q. Yin, R. Cao, X. Q. Gong, T. F. Liu, Journal of Materials Chemistry A 2020, 8, 12370.
